# Structure-based design of a Cortistatin analog with improved immunoregulatory activity against inflammatory bowel disease (IBD)

**DOI:** 10.1101/839787

**Authors:** Álvaro Rol, Toni Todorovski, Pau Martin-Malpartida, Anna Escolà, Elena Gonzalez-Rey, Eric Aragón, Xavier Verdaguer, Mariona Vallès-Miret, Josep Farrera-Sinfreu, Eduard Puig, Jimena Fernández-Carneado, Berta Ponsati, Mario Delgado, Antoni Riera, Maria J. Macias

## Abstract

Ulcerative colitis and Crohn’s disease are inflammatory bowel diseases (IBD) that lead to chronic inflammations of the gastrointestinal tract due to an abnormal response of the immune system. Finding new effective drugs to tackle IBD represents a major therapeutic concern since IBD incidence and prevalence is increasing worldwide. Recent studies positioned Cortistatin (CST) as a candidate for IBD treatment due to its anti-inflammatory and immunomodulatory activity. Here, we studied the structural properties of CST using NMR and synthesized and characterized new analogs displaying enriched populations of some native conformations. One of them, Analog 5, preserved the activity against IBD with an increased half-life in serum, overcoming the native hormone limitation and opening the door for the use of CST analogs as therapeutic agents. This work represents a new approach to the rational design of molecules to treat IBD and a possibility for patients that fail to respond to other therapies.

**Graphical abstract:** 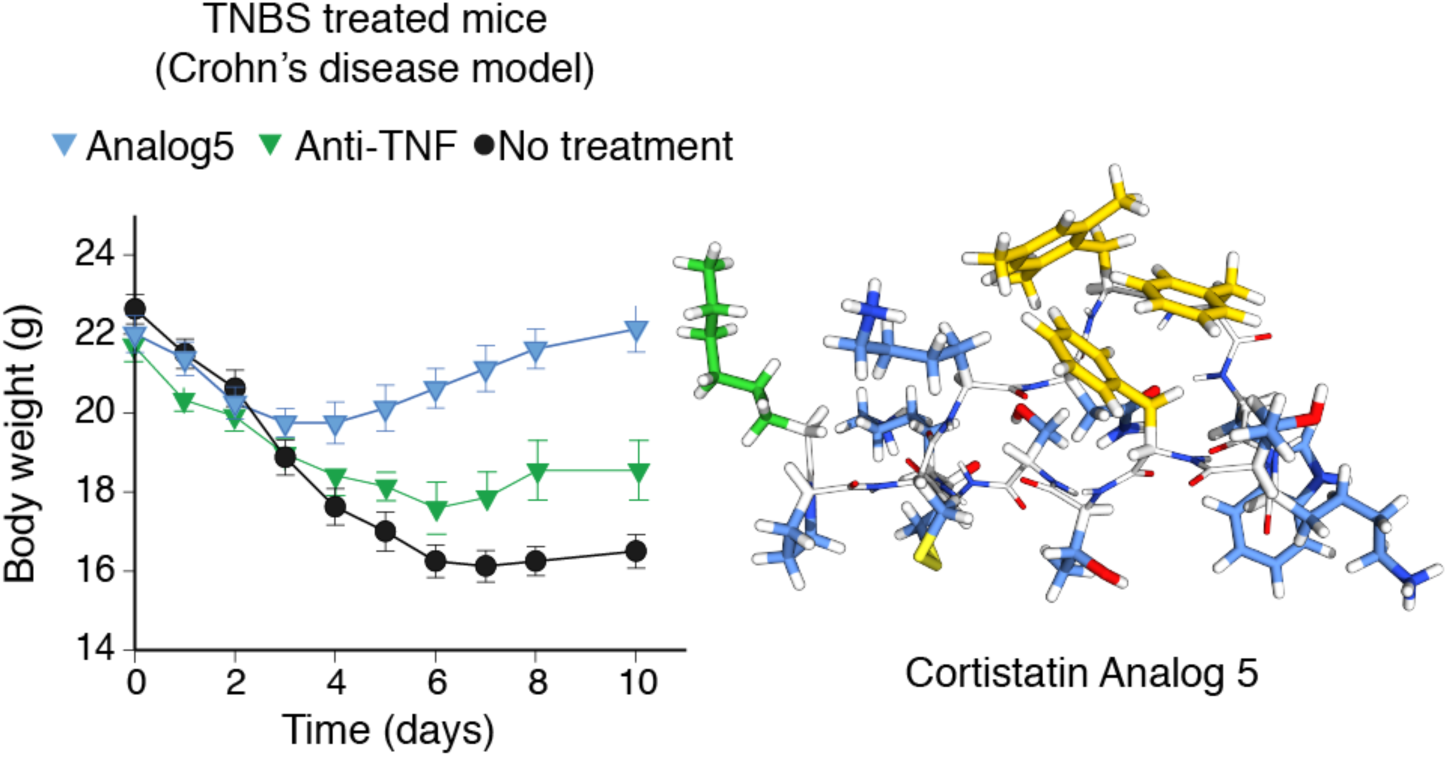

## Introduction

The gastrointestinal tract is a complex system where factors such as diet, environmental and/or microbial inputs challenge the immune system continuously^1^. When these factors dysregulate the gastrointestinal immune system–specially in genetically predisposed individuals –, the perturbation gives rise to inflammation and other symptoms collectively known as Inflammatory Bowel Disease (IBD). In fact, IBD covers a broad spectrum of pathologies and disease severities, including Ulcerative colitis (UC) and Crohn’s disease (CD) that lead to chronic inflammations. The underlying mechanisms involved in these diseases are not fully understood although these mechanisms correlate with elevated levels of pro-inflammatory cytokines, tumor necrosis factor-α (TNF-α), interferon-γ (IFN*γ*) as well as with interleukin 2 (IL-2)^1,2^. In most instances, there is no cure currently available, thus turning IBD into chronic conditions, which often impact the quality of life of the patients. Altogether, these observations position IBD as a major health concern whose treatment is an important priority for our society. Finding specific and safe therapeutic tools to tackle these complex disorders is not trivial and requires an efficient combination of approaches to increase treatment effectiveness, improve patient quality of life and reduce the economic cost^3^.

The current treatment pipeline includes amino-salicylates, corticosteroids, immunomodulators and anti-TNFα antibodies and surgery^4^. Anti-TNF agents have been a major advance as they have proven to be quite effective for both UC and CD but still, one-third of the patients are not responding to the treatment^5^. In this regard, the main challenges that patients face are disease progression, uncontrolled and relapsing inflammation, the appearance of important side-effects in long-term treatments and the high economical cost of them (over $20 billion every year in the USA)^6^. Therefore, finding new molecules to use in combination with, or as alternatives to currently available drugs presents itself as a major need in the field.

Several years ago, cortistatin (CST), a neuro-peptide hormone^7,8^ was found to play immunomodulatory roles^9^ and its external administration protected mice models of IBD from colitis development^10^. In addition, CST mRNA was detected in many neuroendocrine tumors of the lung, and in neuroendocrine tumors of the gastrointestinal tract^11,12^. These evidences place CST as an important signaling molecule of the gastrointestinal track. Moreover, CST was shown to inhibit the in vitro production of inflammatory mediators in activated macrophages, revealing that CST –in contrast to somatostatin–, is an effective agent against hapten-induced CD^11,12^. Cortistatin sequences (human, mouse and rat) are highly similar to that of human somatostatin (SST, Fig. 1a), another natural peptide hormone known to display a broad set of biological actions and a subject of research for many years^13^. These anti-inflammatory effects would be explained by the regulation of cytokines mediated by CST, decreasing the release of pro-inflammatory and cytotoxic cytokines such as TNFα, IFNγ, IL-2, IL-6 or NO, while stimulating the production of anti-inflammatory IL-10. However, the potential application of CST as a therapeutic agent for IBD is limited due to the short lifespan (two minutes in serum) that this peptide exhibits.

**Fig. 1.**
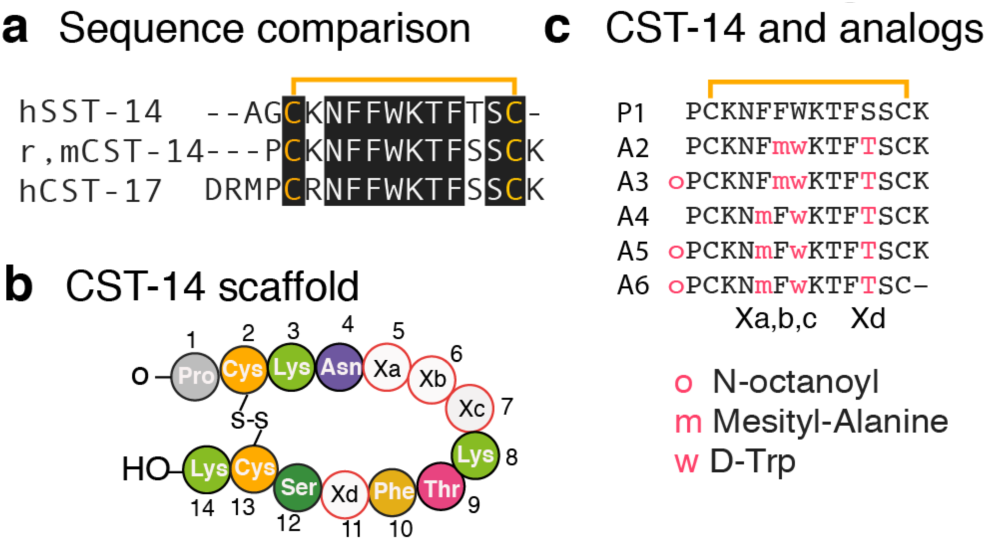
Cortistatin and Somatostatin sequence comparison and analog design. **a.** Sequence comparison of human Somatostatin (SST) to rat and human Cortistatin (CST). **b.** Schematic representation of Cortistatin-14 sequence. Positions Xa, Xb, Xc and Xd have been mutated to prepare the analogs. Native CST was also synthesized for the NMR and functional studies. **c.** Sequences of the analogs prepared in this work. Substitutions are colored. Synthetic scheme is displayed as Supplementary Fig. 1

Despite the sequence similarity of CST and SST, both hormones respond to different stimuli and display different functions. Remarkably, they both bind to the same set of SST receptors (SSTR1-5)^14^. They also share similar limitations in stability, which preclude their direct pharmacological application. In the case of SST, these limitations were overcome by designing new SST analogs containing non-native amino acids and/or by reducing the peptide length. Some of these analogs, such as octreotide (Sandostatin^®^), lanreotide (Somatuline^®^), vapreotide (Sanvar^®^) and pasireotide (Signifor^®^) have successfully reached the market.

Considering that the pharmacophore features of CST are unknown, we contemplate the possibility of improving its stability and immuno-regulatory properties by introducing non-native amino acids at specific possitions –as previously done for SST– while maintaining the full-length peptide chain (Fig. 1b). To assist the rational design of the mutations, we first studied the conformations of the native hormone in solution by nuclear magnetic resonance (NMR). We found that, although the peptide is conformationally flexible, we could observe different aromatic clusters in solution whose conformations were in equilibrium. We hypothesized that these distinct clusters might correlate with the hormone’s function and set to design new analogs that could potentially populate some of these clusters as the main conformation in solution. Among the new peptide analogs designed and tested (Fig. 1c), we observed that analog **5** displayed the anti-inflammatory activity showed by the native peptide in two established preclinical models of IBD^15^. The activity was of the same order of magnitude than that of the anti-TNFα antibody and mesalazine (reference treatments), used for comparison. Moreover, by solving the structure of this analog, we have been able to correlate one specific aromatic cluster with cortistatin’s anti-inflammatory activity.

Our work provides a new framework to design new compounds to treat IBD. These compounds have scafolds based on the natural hormone cortistatin, thus maintaining its immunoregulatory activity while displaying improved stability and specificity with respect to the natural hormone in models of IBD.

## Results

### Conformational properties of Cortistatin in solution by NMR

In contrast to the wealth amount of conformational studies available for somatostatin and its analogues^16-22^, there is no information describing the conformations of cortistatin in solution to date. To bridge this gap, we synthesized cortistatin (peptide **1**, Fig. 1b,c) using standard Fmoc/^*t*^Bu solid-phase peptide synthesis (as described in the methods and in Supplementary Fig. 1)^23^, and studied its folding properties in solution by NMR.

The 2D NMR spectra (experimental section) for this peptide were fully assigned^24^ and the structures were calculated using a manually nuclear Overhauser effects (NOE) assignment protocol in Crystallography & NMR system (CNS)^25^. The pair of conserved cysteine residues enables the peptide to form an internal disulfide bond and a cyclic peptide structure with both N- and C-terminus on one side of the molecule. CST populates an ensemble of conformations in solution, as deduced from the presence of nuclear Overhauser effects (NOEs) between residues separated by more than three residues in the sequence (Fig. 2a). In addition to those NOEs, we also detected additional ones between the side chains of Trp7 and Lys8, denoting the presence of a type II β-turn, at the other side of the disulfide bridge. These same residues and the turn are also observed in SST ^26^. Moreover, weak NOEs were detected between protons of the phenylalanine rings, some of them incompatible with a unique spatial arrangement, confirming the presence of different aromatic clusters in solution. Indeed, the conformational ensemble of CST contains subfamilies of structures mainly defined by the presence of two aromatic clusters, where either two or three aromatic Phe residues pack together (Fig. 2b-d). Overall these results suggest that different sets of conformations can be adopted by the native hormone in solution, and perhaps, in a context dependent environment, these different orientations might be specifically selected by distinct protein/receptor partners in order to regulate cellular responses.

**Fig. 2.**
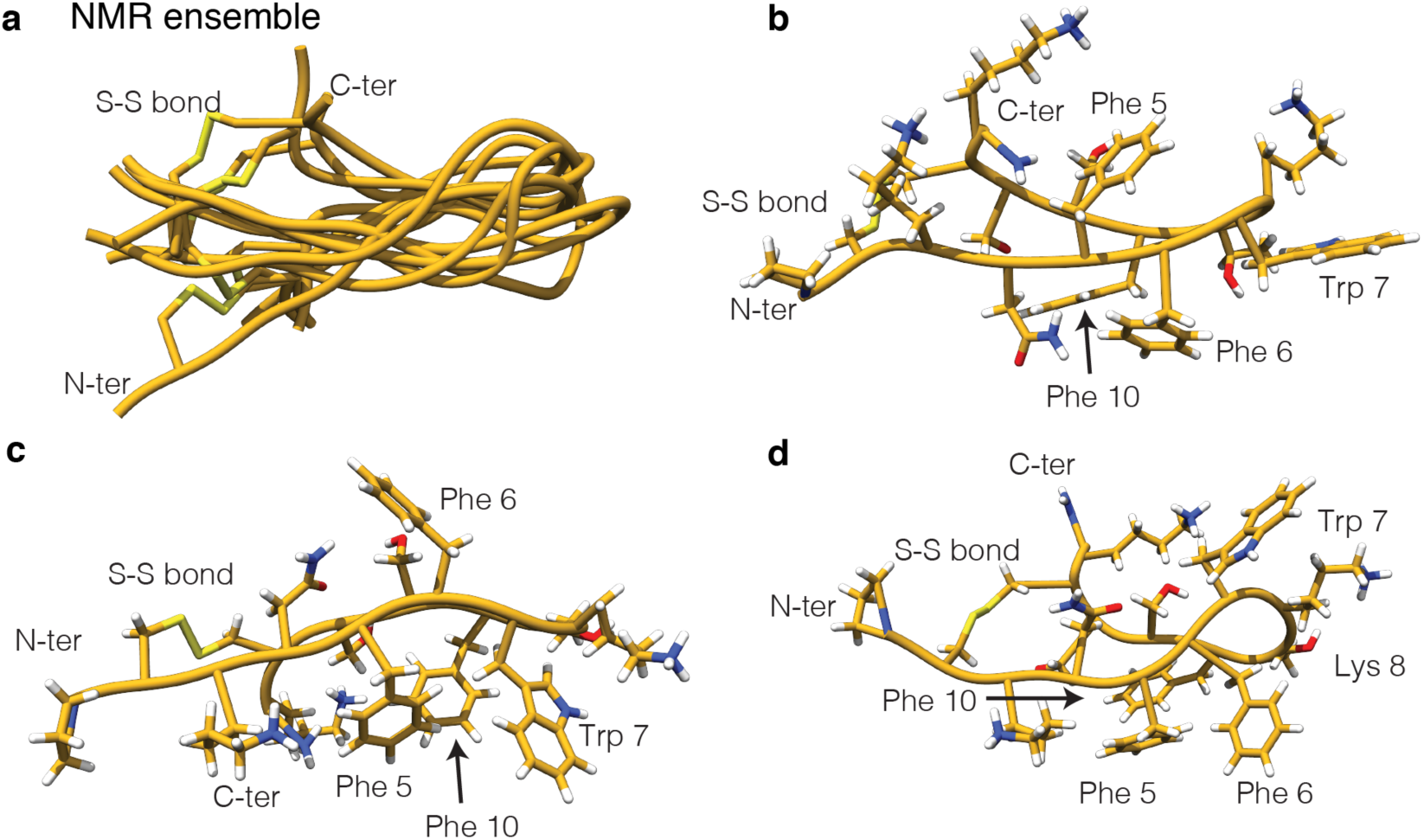
Cortistatin NMR structures. **a.** Backbone superposition of the 10-lowest energy structures. The peptide backbone is shown in gold and the N- and C-terminus residues as well as the disulfide bond are labeled. **b-d**. Representative aromatic clusters. All side chains are displayed to facilitate the structural analysis. Side chains participating in the interactions are specifically labeled.

### *De novo* design of cortistatin analogs

Encouraged by these results we designed new CST analogs, envisioning the possibility that they could individually populate –as the main conformation– each of these aromatic clusters (Fig. 1b,c, analogs **2-6**). We also aimed to correlate the conformations present in the native CST-14 ensemble and in the analogs, with a given observable in our functional assays, enabling the future design of new molecules with immunoregulatory activity and improved stability.

In order to design the first CST analogs, we collected the information described in the literature to optimize the stability and properties of SST. For instance, it is well known that substitution of L-Trp to D-Trp increases the population of the native β-turn, as well as the hormone activity and its half-life in serum^27^. It has also been shown that substitutions of Phe, by L-mesityl alanine (3,5-dimethylphenylalanine, Msa) favored π-π interactions between aromatic rings and increased the overall conformational stability of SST derivatives^19,20^. For these reasons, in all CST analogs L-Trp was replaced by D-Trp in position 7 and Ser11 was replaced by Thr11 to stabilize the β-turn; and either Phe6 or Phe5 were replaced by Msa (analogs **2** and **4**) (Fig. 1b,c). In addition to these modifications, terminal residues were acylated as N-octanoyl amides (analogs **3** and **5**). The logic for this modification was to increase the hydrophobicity of the analog and to mimic a similar modification observed in Ghrelin, a hormone produced in endocrine cells in the digestive tracts and that shares with CST the ability to interact with the growth hormone secretagogue receptor GHSR-1a ^28^. Finally, a deletion of L-Lys14 (analog **6**) was synthesized with the aim of improving the stability of the analogs, as this deletion was identified as the main degradation product of the native hormone by Mass Spectrometry.

All analogs were prepared by standard Fmoc/^*t*^Bu solid-phase peptide synthesis (SPPS) as the wild type CST peptide. The Fmoc-protected Msa was synthetized by asymmetric hydrogenation following our previously described protocol^29^. For the NMR structural studies, the peptides were used as TFA salt and for the *in vitro* and *in vivo* studies the TFA counter ion was replaced by acetate salts using ion exchange chromatography.

### Half-life in serum and *in vitro* activity of the new analogs

The stability of the natural CST-14 hormone and of the new analogs was evaluated measuring their half-life in serum and their *in vitro* affinity by analyzing their binding ability to Somatostatin receptors. Modification of the native cortistatin peptide significantly increased its half-life in serum going from 2 min (cortistatin) to 21 min and 2100 min for analogs **5** and **6** respectively. These results indicated that substitutions introduced in these analogs resulted in at least a 10-fold increase of stability with respect to the native neuropeptide. In fact, the main metabolite of peptide **5** is peptide **6**. Unfortunately, the lack of Lys14 in analog **6**, also reduced its aqueous solubility, limiting its use in our functional and structural assays.

Since cortistatin binds *in vitro* with nanomolar affinity to the five known Somatostatin receptors, we set to investigate if the new analogs maintained similar profiles^19,20,22,30,31^. Our results corroborate these findings and reveal that both native hormones and analogs **2-3** have similar profiles and a high preference towards receptor 2, as all SST analogs currently in the market. Remarkably, analogs **4** and **5** have distinct binding preferences for SST receptors (Table 1). Analog **5** exhibits a binding preference for receptors 3 and 5. Analog **5** binding to receptor 4 is close to that of SST, and binding to receptor 2 is much lower when compared to both native CST and SST hormones.

**Table 1.** Pharmacological profile of Cortistatin, Somatostatin and analogs 2-5.

|  | SSTR1 (nM) | SSTR2 (nM) | SSTR3 (nM) | SSTR4 (nM) | SSTR5 (nM) |
| --- | --- | --- | --- | --- | --- |
| **Somatostatin** | 1.9 | 0.016 | 0.25 | 1.55 | 0.76 |
| **Cortistatin** | 7.5 | 0.091 | 2.72 | 7.54 | 3.48 |
| **Analog 2 (A2)** | 13.4 | 0.26 | 5.1 | 65 | 14.9 |
| **Analog 3 (A3)** | 140 | 0.30 | 32 | 210 | >100 |
| **Analog 4 (A4)** | 11.2 | 4.33 | **0.62** | 8.59 | 2.58 |
| **Analog 5 (A5)** | 25.0 | 16 | **0.91** | 2.39 | **0.53** |

The pharmacological profile (K<sub>a</sub>, nM) of Cortistatin, Somatostatin and analogs 2-5 against five Somatostatin receptors. Experiments were performed as duplicates. Similar values to those of native Somatostatin and Cortistatin are highlighted in bold.

We then moved to evaluate the potential immunosuppressive and anti-inflammatory effects of our peptides in two *in vitro* assays previously established^15,32-34^. Analogs **2, 3** and **4** showed no effect on the inflammatory responses in activated lymphocytes and macrophages (Table 2). Conversely, CST and its analog **5** strikingly reduced the production of both the inflammatory mediators TNFα, IL-6 and nitric oxide (NO) by murine macrophages activated with a bacterial endotoxin (lipopolysaccharide, LPS) and reducing the proliferative response and the production of cytokines IFNγ and IL-2 by activated mouse spleen cells as well (Table 2).

**Table 2.** Effects on lymphocyte and macrophage activation.

| Peptides | Splenocytes<sup>a</sup> |  |  | Macrophages<sup>b</sup> |  |  |
| --- | --- | --- | --- | --- | --- | --- |
| | Proliferation<br>(cpm) | IFNγ<br>(ng/mL) | IL-2<br>(ng/ml) | TNFα<br>(ng/mL) | IL-6<br>(ng/mL) | NO<br>(nmol/L) |
| **Unstimulated** | 640 | 0 | 640 | 0.48 | 0 | 0.44 |
| **Activated** | 9843 | 2.52 | 9843 | 5.78 | 4.48 | 5.23 |
| **+Somatostatin** | 9936 | 2.31 | 9936 | 5.21 | 4.20 | 4.32 |
| **+Cortistatin** | 5500 | 1.13 | 5500 | 3.17 | 3.78 | 3.08 |
| **+Analog 2 (A2)** | 9866 | 2.52 | 9866 | 4.90 | 4.23 | 4.43 |
| **+Analog 3 (A3)** | 9316 | 2.53 | 9316 | 3.84 | 3.83 | 4.49 |
| **+Analog 4 (A4)** | 10090 | 2.56 | 10090 | 4.04 | 3.79 | 4.36 |
| **+Analog 5 (A5)** | **5863** | **1.33** | **5863** | **2.51** | **2.77** | **3.04** |

<sup>a</sup>Mouse spleen cells were stimulated with anti-CD3 antibodies (2 $\mu$ g/mL) in the absence or presence of 100 nmol/L of peptides. Cells incubated with medium alone were used as unstimulated controls. After 48 h, IL-2 and IFNγ levels in the supernatants were determined by ELISA. After 72 h, [<sup>3</sup>H]-thymidine was added to cultures and proliferation was determined 8 h later by incorporation of thymidine to DNA. Results show mean of three independent cell cultures.
<sup>b</sup>Raw 264 macrophages were stimulated with LPS (1 $\mu$ g/mL) in the absence or presence of 100 nmol/L of peptides. Cells incubated with medium alone were used as unstimulated controls. After 24h, TNFα and IL-6 levels in the supernatants were determined by ELISA, and nitric oxide levels in the supernatants were assayed by nitrite contents by using a Griess reaction. Results show mean of three independent cell cultures.
Analog 5 values are highlighted in bold for comparison.

Our results indicate that analog **5** showed the best results both in anti-inflammatory and immunosuppressive assays as well as in stability in serum and displayed a preference for receptors 3, 4 and 5, different from native SST and CST hormones that preferentially select receptor 2.

### Analog 5 as a promising candidate in the treatment of IBD

Once the immunosuppressive activity of analog **5** in vitro was confirmed, we next investigated its potential therapeutic action in two established experimental models of acute and chronic colitis induced by oral administration of dextran sulfate sodium (DSS) and by intrarectal infusion of 2,4,6-trinitrobenzene sulfonic acid (TNBS), which displays clinical, histopathological and immunological features of human UC and CD, respectively ^32,33,35^.

In the DSS model, intestinal inflammation results from the impairment of the intestinal epithelial cell barrier function by DSS, subsequent exposition of the submucosa to various luminal antigens (bacteria and food) and activation of the inflammatory cells involved in the innate immunity. By 7 days, oral administration of 5% DSS resulted in a progressive increase in the disease activity index, characterised by acute colitis, bloody diarrhoea and sustained weight loss resulting in 50% mortality (Fig. 3a). Colitis was correlated with dramatic signs of colon damage and inflammation, characterised by colon shortening and increased organ weight (Fig. 3a). Systemic treatment with analog **5** during three days avoided mortality, ameliorated body weight loss, and improved the wasting disease and colon inflammation, similarly to native cortistatin (Fig. 3a).

**Fig. 3.**
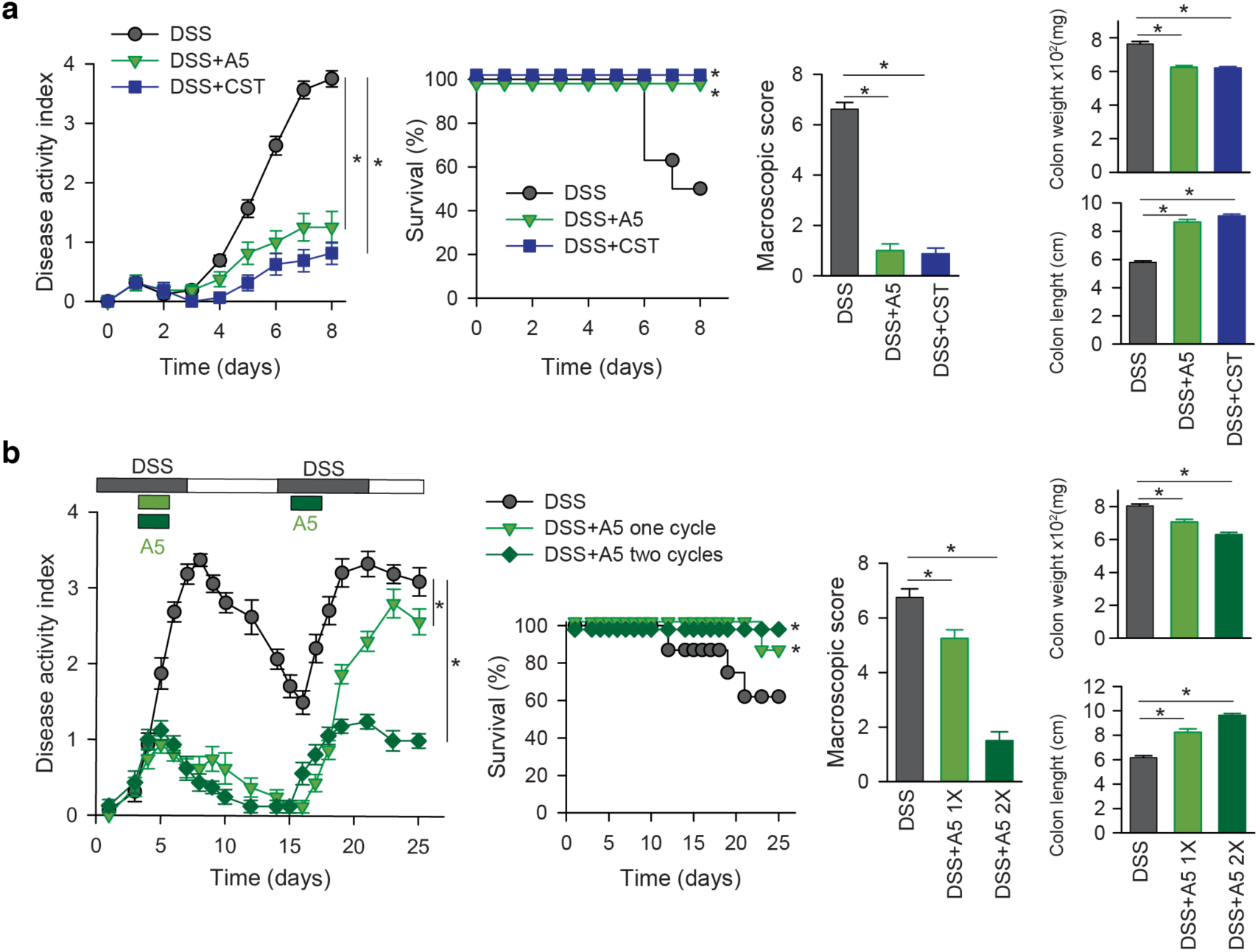
Therapeutic effect of analog 5 in DSS-induced ulcerative colitis. **a.** *Analog 5 protects from acute ulcerative colitis.* Mice received 5% DSS with their drinking water from day 0 to day 7. Cortistatin (CST) or analog 5 (A5) were administered intraperitoneally at a dose of 0.4 mg/kg on days 1, 2 and 3 after initiation of DSS treatment. Disease activity indexes (scoring body weight loss, stool consistency and presence of fecal blood, scale 0-4) and survival rate were daily evaluated. Signs of colon inflammation were determined at day 8 by measuring macroscopic damage scores (scale 0-8) and colon length (in cm) and weight (in mg). **b.** *Protective effect of A5 in chronic ulcerative colitis.* Mice received 3% DSS with their drinking water in a cyclic manner. Each cycle consisted of 7 days of DSS followed by a 7-day period without DSS supplementation. Analog 5 (0.4 mg/kg) was injected subcutaneously during the first cycle of DSS on days 3 to 5 (1X), or during the two cycles of DSS on days 3 to 5 and on days 16 to 18 (2X). Disease activity scores and mortality were determined daily and sings of colon inflammation were measured at day 25 or immediately after death of the animal. Mice receiving tap water instead of DSS were used as naïve controls and showed no clinical signs (see supplementary Fig. S3). n=8 mice/group. *p<0.05 versus untreated DSS-colitic mice.

Treatment with analog **5** was also effective in a remitting-recurrent model of ulcerative colitis induced by administration of 3% DSS in two cycles. Subcutaneous injection of analog **5** during the first acute peak of colitis significantly reduced the clinical activity, as shown by improvement of stool consistency, less rectal bleeding, decrease in colon shortening, amelioration of colon damage and improvement in survival rate, and conferred significant resistance to disease activity during a second cycle of DSS administration (Fig. 3b). A second injection of analog **5** during the second cycle of DSS infusion almost completely abrogated the clinical signs (Fig. 3b). Remarkably, treatment with analog **5** was even more efficient than a therapy of reference based on a neutralizing anti-TNFα antibody (Supplementary Fig. 3).

In the TNBS model, intestinal inflammation results from initial rupture of intestinal barrier by 50% ethanol and later TNBS-mediated haptenization of autologous host mucosal proteins and subsequent stimulation of a Th1 cell-mediated immune response against TNBS-modified self-antigens. The systemic injection of analog **5** during the progression of TNBS-induced acute colitis significantly protected from development of the profound and continuous body weight loss, bloody diarrhea and mortality caused by severe colonic inflammation (Fig. 4a). Interestingly from a therapeutic point of view, treatment with analog **5** in a curative regimen at the peak of the disease also resulted significantly effective reducing clinical signs and improving survival (Fig. 4b), showing even better effects than those with reference treatments, such as a neutralizing TNF therapy or oral administration of mesalazine (Supplementary Fig. 4a). We observed that both intraperitoneal and subcutaneous administration pathways showed similar effectiveness whereas oral administration was less effective ameliorating TNBS-induced acute colitis (Fig. 4b). Interestingly, in a model of chronic colitis induced by repetitive weekly infusion of low doses of TNBS, an initial injection of analog **5** during the first week significantly protected from subsequent expositions to TNBS (Fig. 4c), suggesting the induction of a kind of tolerance to disease recurrence. However, treatment with mesalazine or with anti-TNFα antibody fully or partially failed to induce this tolerance effect (Supplementary Fig. 4b). As expected, repetitive treatments with analog **5** after each TNBS infusion almost completely reversed disease evolution (Fig. 4c), in a similar way than that resulted from chronic administration of mesalazine or repetitive injections of anti-TNFα antibody (Supplementary Fig. 4b).

**Fig. 4.**
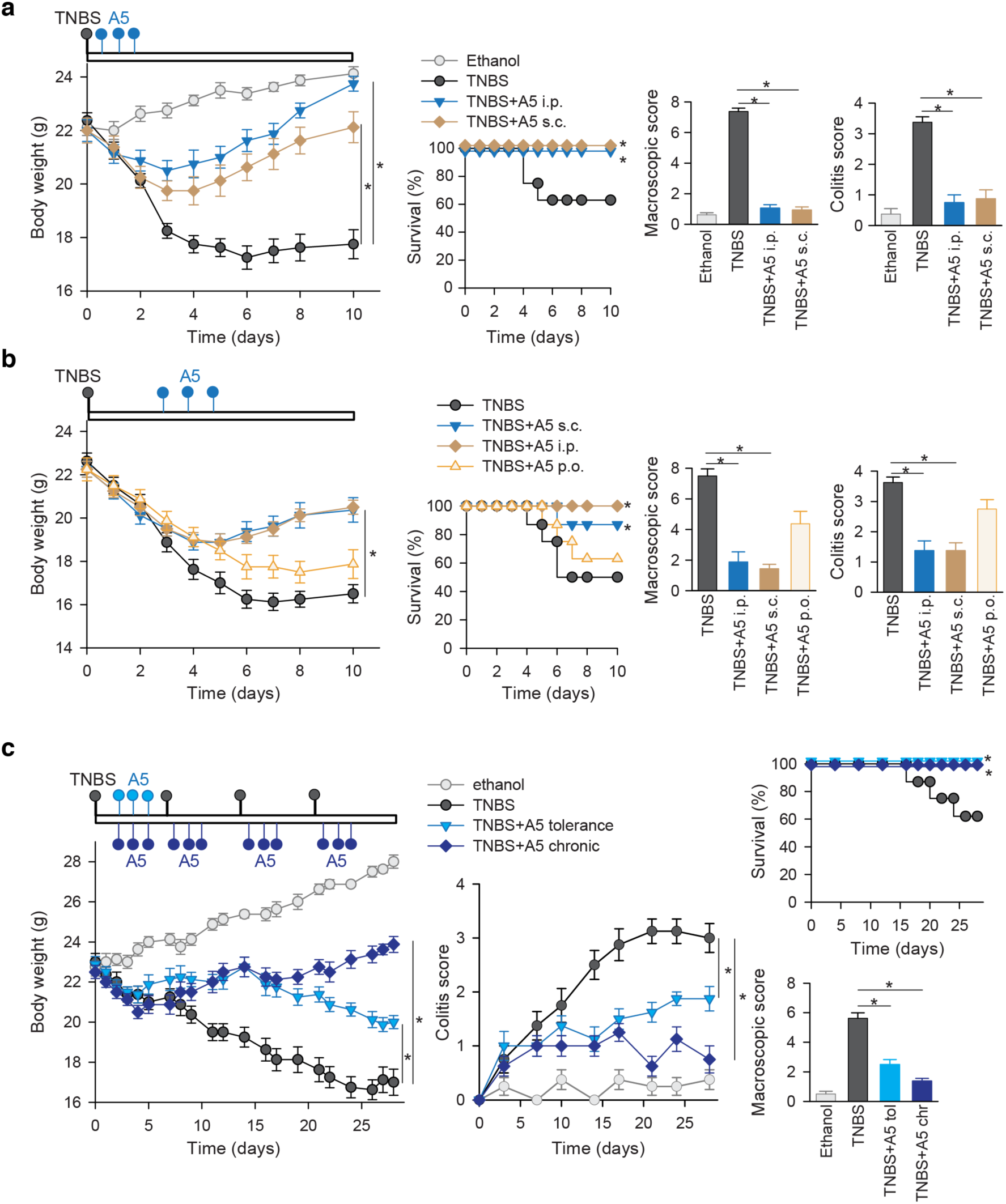
Therapeutic effects of analog 5 on TNBS-induced acute and chronic colitis. **a and b.** *Analog 5 protects from TNBS-induced acute colitis.* Acute colitis was induced in male Balb/c mice by intrarectal administration of TNBS (3 mg/mouse) in 50% ethanol. Mice were treated intraperitoneally (i.p.), subcutaneously (s.c.) or orally (p.o.) three times with analog 5 (A5, 0.4 mg/kg) following a protective regime at 12, 24 and 48h after TNBS infusion (a) or a curative regime 3, 4 and 5 days after TNBS infusion (b). Mice injected intrarectally with 50% ethanol were used as basal controls. Disease evolution and severity was monitored by survival and weight loss (expressed in grams). Colitis score (scale 0-4) was determined at day 4 (protective regime) or at day 6 (curative regime). Macroscopic damage score (scale 0-10) was evaluated in colons isolated at day 10 or immediately after death of animal. **c.** *Therapeutic effect of analog 5 in TNBS-induced chronic colitis.* Mice were injected intrarectally once a week with increasing doses of TNBS (0.8 mg/mouse at day 0, 1 mg/mouse at day 7, 1.2 mg/mouse at day 14 and 1.5 mg/mouse at day 21). Analog 5 was injected s.c. at 0.4 mg/kg in saline, following a tolerance regime at days 3, 4 and 5 after TNBS injection or a chronic treatment at days 3, 4, 5, 8, 9, 10, 15, 16, 17, 22, 23 and 24. Mice injected with 50% ethanol on days 1, 7, 14 and 21 were used as basal controls. Disease evolution and severity were evaluated by determining body weight loss, survival, colitis score and the macroscopic colonic damage (at day 28 or after death of each animal). n=8 mice/group. *p<0.05 versus untreated TNBS-colitic mice.

Overall, these results in animal models position analog **5** as a potential candidate in the treatment of IBD.

### Conformational studies of Cortistatin and its analogs by NMR

With the aim to understanding the different activity profiles of these analogs we have analysed their conformational properties by NMR, starting with analog **5**, which is the most biologically active of all. The NMR data allowed us to characterize a major conformation in solution due to well dispersed signals in the 2D TOCSY and to the presence of abundant and unambiguous NOEs in the 2D NOESY spectra. The assignment of the 2D TOCSY and NOESY experiments indicates that Msa5, Phe6 and Phe10 aromatic rings are clustered on one side of the peptide plane, with the rings of Msa5 and Phe10 oriented edge-to-face, and with the aromatic rings of Phe6 and 10 arranged offset-stacked, all pointing away from the D-Trp7:Lys8 pair (Fig. 5a, Supplementary Fig. 5). These NOEs were also present in the native SST hormone **1** but are more intense in analog **5** (Fig. 2c,d), suggesting that this analog displays an enriched set of conformations already present in the pool of conformations sampled by the native hormone.

**Fig. 5.**
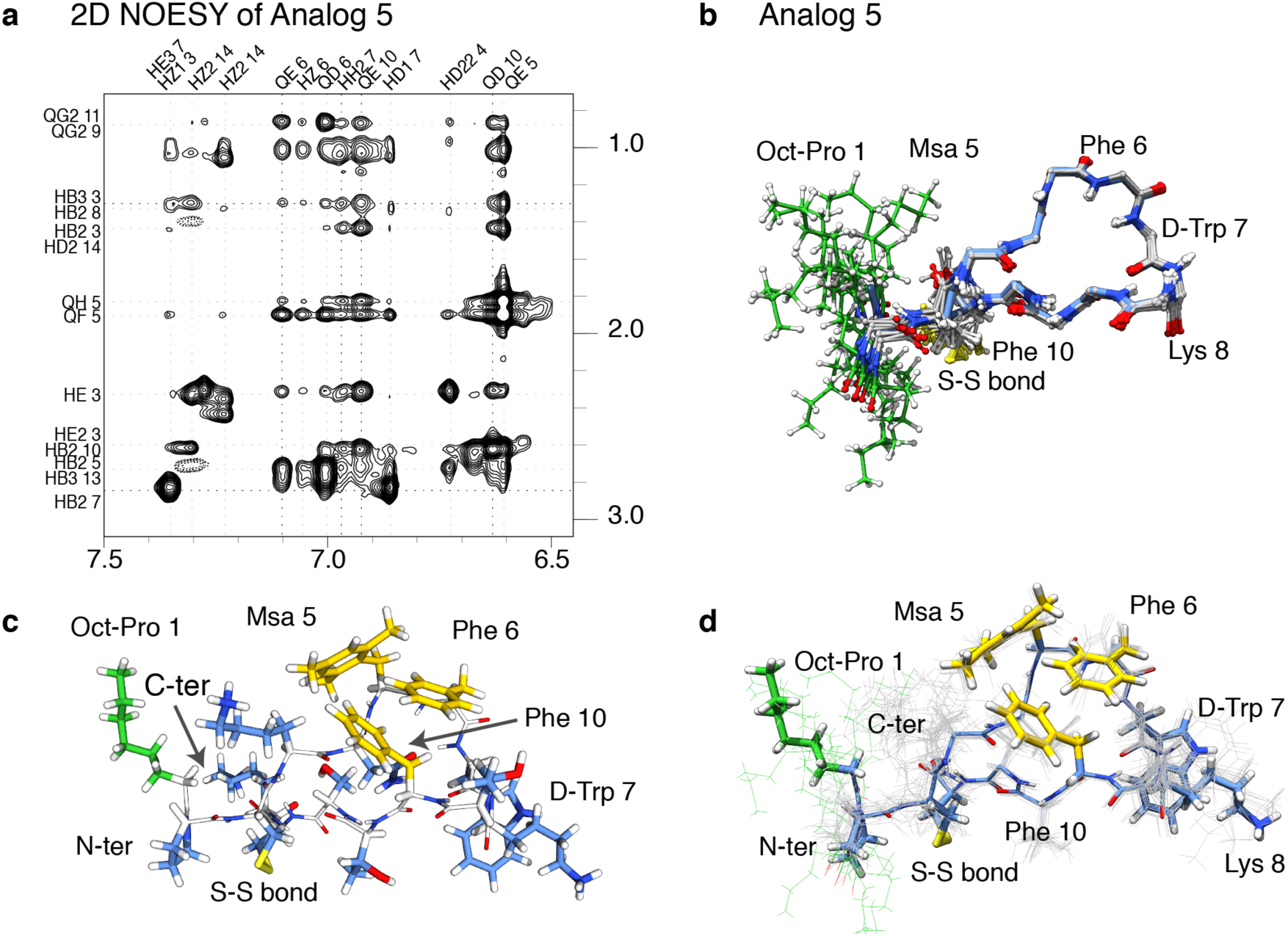
NMR structure of analog 5. NMR restraints and ensemble of structures corresponding to analog **5. a.** Sidechain amide region of the analog **5**(^1^H-Noesy experiment), with peak assignments. **b.** Backbone superposition of the 15 best calculated structures. **c.** Analog **5**, lowest energy structure. Side chains are depicted as sticks. The residues defining the aromatic cluster are shown in yellow, the octanoyl moiety is highlighted in green whereas the remaining residues are colored in blue. The peptide backbone is shown in white. **d.** Sidechain and backbone distribution for the ensemble of 15 best structures. Some residues are labeled.

The backbone superposition of the 15 best structures (according to structure validation tests and energy values) is shown as Fig. 5b whereas the side chain orientation of some selected amino acids and all sidechains are shown in Fig. 5c,d. The orientation of the octanoyl moiety is not restrained in the calculation since only few and weak NOEs were observed from the octanoyl residue to Lys4/12 side chains. Remarkably, it seems that both Lys4 and Lys12 side chains shield the octanoyl moiety from the rest of the peptide surface (Fig. 5a), defining a hydrophobic pole adjacent to the aromatic cluster located in the middle of the structure. These hydrophobic areas do not prevent the peptide from being soluble but might enhance the interaction with CST natural binders/receptors in a cellular context, explaining why this analog is more active than analog **4**, (identical in sequence to **5** but lacking the octanoyl moiety).

We also characterized the conformational properties of analogs **2, 3** and **4.** NMR data acquired for them revealed a pattern of NOEs similar for these three analogs but different from analog **5.** In these cases (Fig. 6a,c,e), the aromatic residues 5 and 10 aligned on one side of the molecule, whereas the side chain of residue 6 is located on the other side of the peptide plane. In this conformation, D-Trp7 is close to the pair of aromatic rings 5 and 10, due to the presence of weak NOEs involving the ring protons of D-Trp7 and the methyl protons of Msa5. Among the three analogs, analog **3** that also contains an octanoyl moiety and the Msa at position 6 is less rigid than analogs **2** and **4** (Fig. 6b,d,f). Moreover, in analog **3**, the reduced number of NOEs between the pair D-Trp7:Lys8 also corroborate the presence of conformational variability in this part of the structure.

**Fig. 6.**
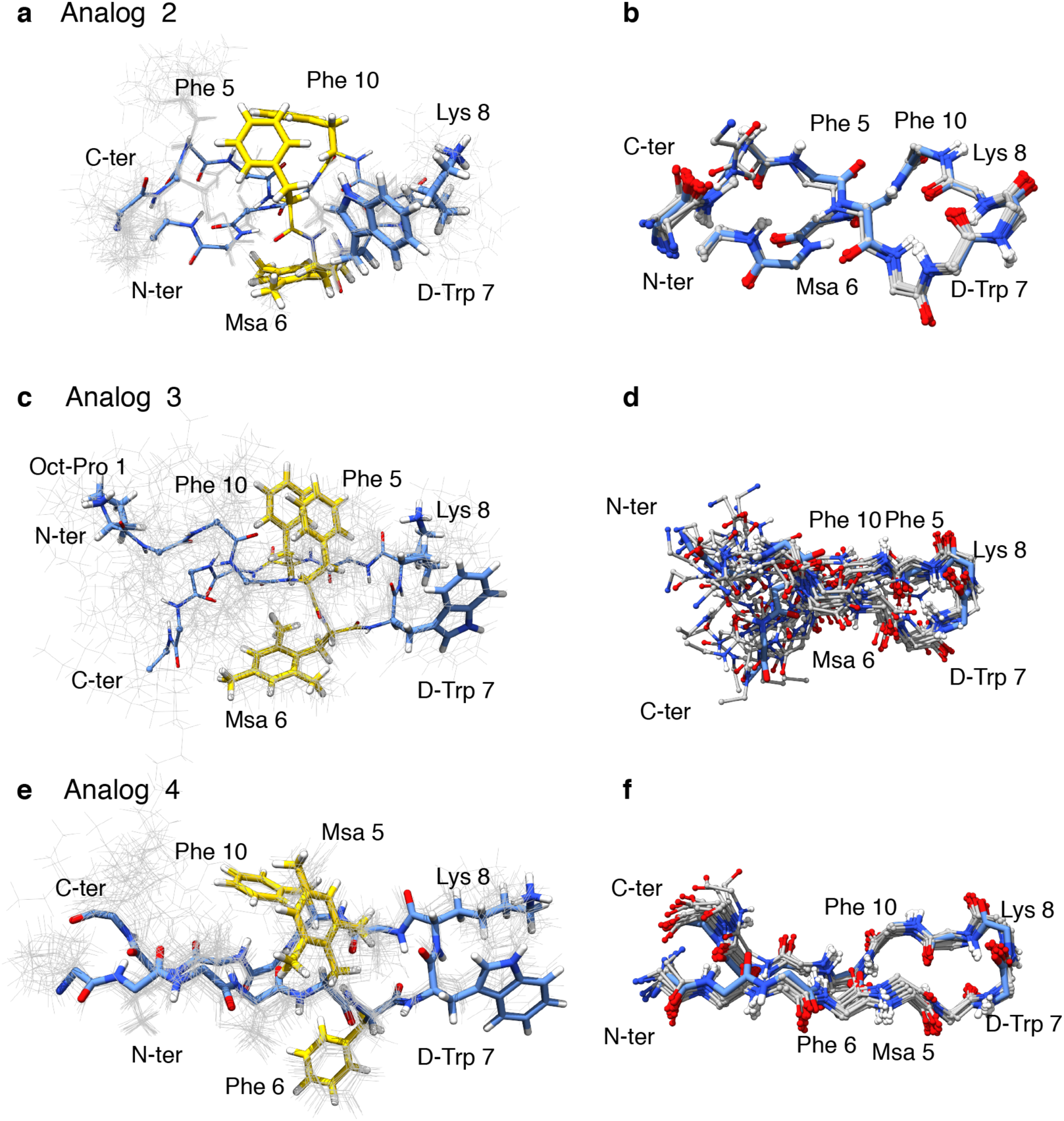
NMR structures corresponding to analogs 2, 3 and 4. **a.** Analog **2**, sidechain and backbone distribution for the ensemble of 15 best structures. Side chains are depicted and labeled. The lowest energy structure is highlighted as sticks. **b.** Backbone superposition of the 15 best calculated structures. Some positions are indicated. **c-f** Same panels are displayed for analogs **3** and **4**.

Finally, we found that the loss of Lys14 (analog **6**) had a significant impact in the solubility of the sample, preventing the acquisition of 2D NMR experiments in aqueous buffer solution and explaining its poor activity in the cellular assays.

All in all, our studies indicate that the combination of Msa in position 5 and of the octanoyl moiety at the N-terminus contribute to favor the cluster formation observed in analog **5**, since the other analogs, either lacking such octanoyl moiety (analog **4**) or with the Msa modification at a different position (analog **2** and **3**), do not adopt this structural arrangement.

## Discussion and Conclusions

IBD is an emerging global disease and its impact is increasing in children and adolescents. Its rapid development runs along with demographic shifts (urbanization) and an impact of environmental factors (diet, microbiota and psychological stress).

Cortistatin is expressed in immune cells and it has been shown to be a potent anti-inflammatory agent that can deactivate some of the IBD responses in animal models^10^. However, the native sequence has other roles in modulating the GH-releasing activity of Ghrelin and parallels some roles of SST. We have expanded the repertoire of molecules that can help treating patients, designing analogs that retain the anti-inflammatory capacity of the hormone, with increased stability. These new analogs were the result of introducing native and non-native amino acids and other modifications into the native scaffold. We found that analog **5** displayed a ten-fold stability increase in serum while preserving the pattern of immunosuppressive and anti-inflammatory activity of cortistatin. This analog **5** contains a Msa in position 5 and an octanoyl moiety at the N-terminus as main modifications of the native sequence.

With respect to binding to SST receptors, cortistatin is known to interact with all of them in the low nanomolar range^7^. Our results corroborate these findings and reveal that analog **5** has a distinct binding preference for SST receptors, selecting receptors 3, 4 and 5 instead of receptor 2. This observation opens the possibility of using this analog to inhibit prolactin secretion in prolactin-secreting adenomas since these roles are ascribed to specific functions of the SST receptor 5^36,37^. Certainly, additional experiments are required to test this hypothesis.

The immunosuppressive and anti-inflammatory activities of the analogs were measured using cytokines and inflammatory factors in vitro and corroborated by experiments performed in mouse models of IBD. Because analog **5** mimicked the immunosuppressive activity of cortistatin in vitro, it is plausible that the effect observed for analog **5** in experimental colitis is exerted by regulating the two pathological components of IBD, i.e., exacerbated mucosal inflammation and Th1-driven self-reactive responses, as previously described by cortistatin^9,10,38^. Our results revealed that treatment with analog **5** was more effective than the administration of the anti-TNFα antibody used as the reference treatment in these assays, highlighting the potential of this new molecule.

To rationalize the observed biological properties, the new peptides were characterized using NMR, results showing that these analogs populate two main set of conformations in solution. We found that the most active peptide, analog **5**, displays an aromatic cluster in solution, defined by contacts between the aromatic rings of Phe6, Phe10 and Msa5 absent in the other analogs. We suggest that receptors or binding partners present in inflammatory cells might recognize epitopes similar to the aromatic cluster found in this analog **5**. We believe that our work opens a new path to the rational drug design to help control inflammatory bowel syndromes, using a new family of compounds derived from natural hormones.

## Author contribution

M.D., A.R., B.P. and M.J.M. designed and supervised the project. A. Rol, P.M-M.; E.A. and M.J.M. assigned and analyzed the NMR data, performed NMR measurements and computational analysis. A. Rol, A.E., T.T., and M.V-M synthesized the peptides. J.F-C, J.F-S supervised the synthesis. E.G-R and M.D. performed mouse experiments. E.P assisted the first and senior authors with manuscript coordination. M.J.M. wrote the paper with input from P.M-M.; M.D.; J.F-C and A.R. All authors contributed ideas to the project.

## Acknowledgements

A. Rol was a recipient of a PhD fellowship granted by the Generalitat de Catalunya (FI) and A.E. and E.P. were recipients of PhD fellowships granted by the Severo Ochoa Program (FPI). T.T. was a postdoctoral fellow co-funded by the Marie Skłodowska-Curie COFUND actions (IRB Barcelona Interdisciplinary Postdoc Programme). This work was supported by by the the Spanish Ministry of Economy, Industry and Competitiveness (MINECO) grants: CTQ2014-56361-P and CTQ2017-87840-P (A.R.) and by BCN PEPTIDES S.A. We also acknowledge institutional funding from the Spanish Ministry of Economy, Industry and Competitiveness (MINECO) through the Centres of Excellence Severo Ochoa award given to the IRB Barcelona as well as from the CERCA Program of the Catalan Government. MJ.M. is an ICREA Programme Investigator.

## Conflicts of interest

Analog 5 is patented, EP 3046933 B1 (BCN PEPTIDES, S.A.) 2019-02-27, “Cortistatin analogues for the treatment of inflammatory and/or immune diseases”.

Methods section is provided as Supplementary information.

## Figures

**Supplementary Fig. 1.**
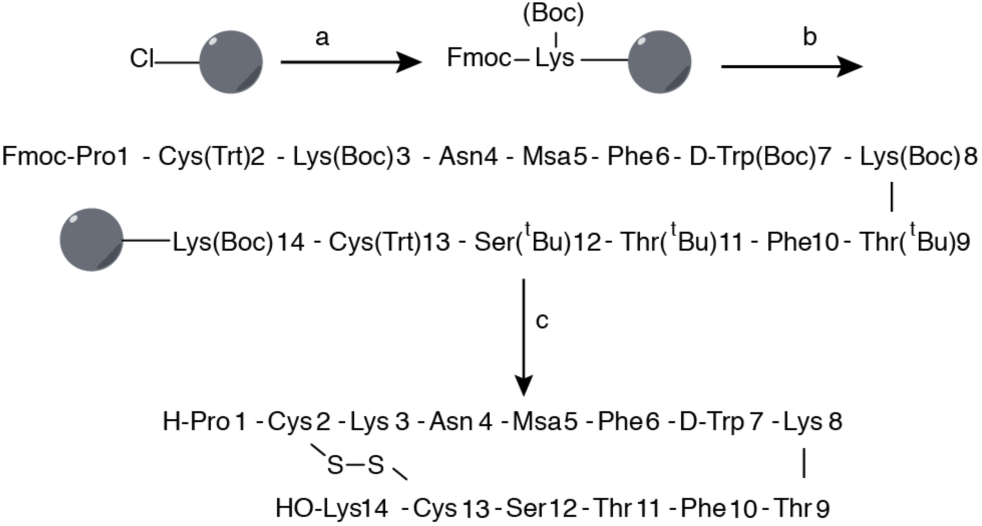
Synthetic scheme. Synthesis of [L-Msa5_D-Trp7_L- Thr11]-CST14(analog 4). All peptides have been prepared following a similar approach. a) 1. Fmoc-L-Lys(Boc)-OH (3 equiv) or Fmoc-L-Cys(Trt)-OH (3 equiv), DIEA (3 equiv), 2. MeOH; b) 1. Piperidine 20% DMF, 2. Fmoc-Aaa-OH (1.5–3 equiv, when the Msa amino acid was coupled, only 1.5 equivalents were used), DIPCDI (3 equiv), HOBt (3 equiv), DMF (x12), 3. Piperidine 20% DMF, 4. Fmoc-Pro-OH, DIPCDI, HOBt, DMF c) 1. CH_2_Cl_2_/TFE/AcOH 2. I_2_, 3. TFA/CH_2_Cl_2_/anisole/H2O. Boc=*tert*-butoxycarbonyl, DIEA=diisopropylethylamine, PCDI=diisopropylcarbodiimide, DMF= N,N’-dimethylformamide, Fmoc=N-(9- fluorenylmethoxycarbonyl), HOBT=1-hydroxybenzotriazole, TFA=trifluoroacetic acid, TFE=2,2,2-trifluoroethanol.

**Supplementary Fig. 3.**
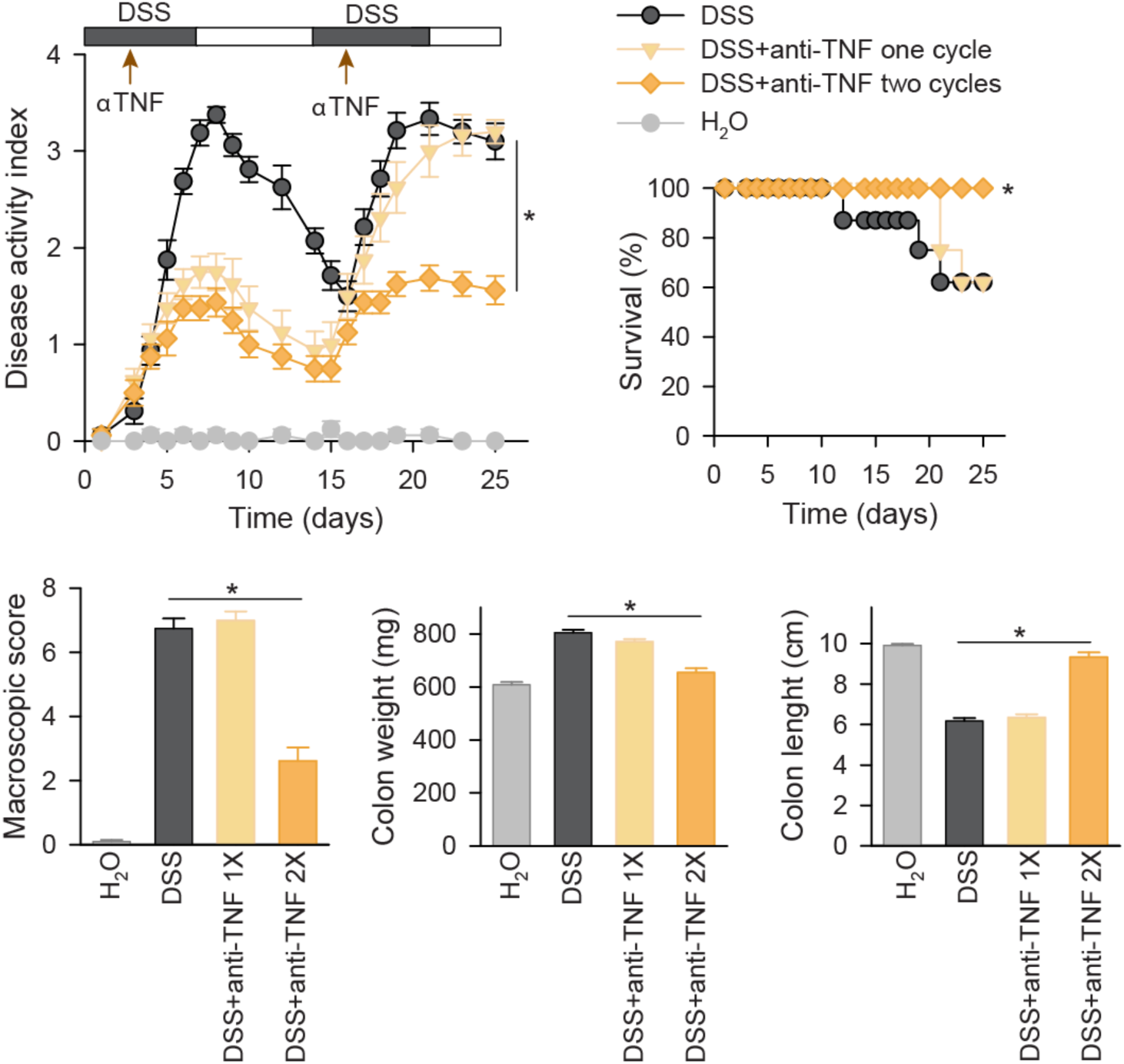
Therapeutic effect of anti-TNFα antibodies in DSS-induced chronic ulcerative colitis. Mice received 3% DSS with their drinking water in a cyclic manner. Each cycle consisted of 7 days of DSS followed by a 7-day period without DSS supplementation. Anti-mouse TNFα antibody (5 mg/kg) was injected intravenously during the first cycle of DSS at day 3 (1X), or during the two cycles of DSS at days 3 and 16 (2X). Disease activity indexes (scoring body weight loss, stool consistency and presence of fecal blood, scale 0-4) and survival rate were daily evaluated. Signs of colon inflammation were determined at day 25 or immediately after death of each animal by measuring macroscopic damage scores (scale 0-8) and colon length (in cm) and weight (in mg). Mice receiving tap water instead of DSS were used as naïve controls. n=8 mice/group. *p<0.05 versus untreated DSS-colitic mice.

**Supplementary Fig. 4.**
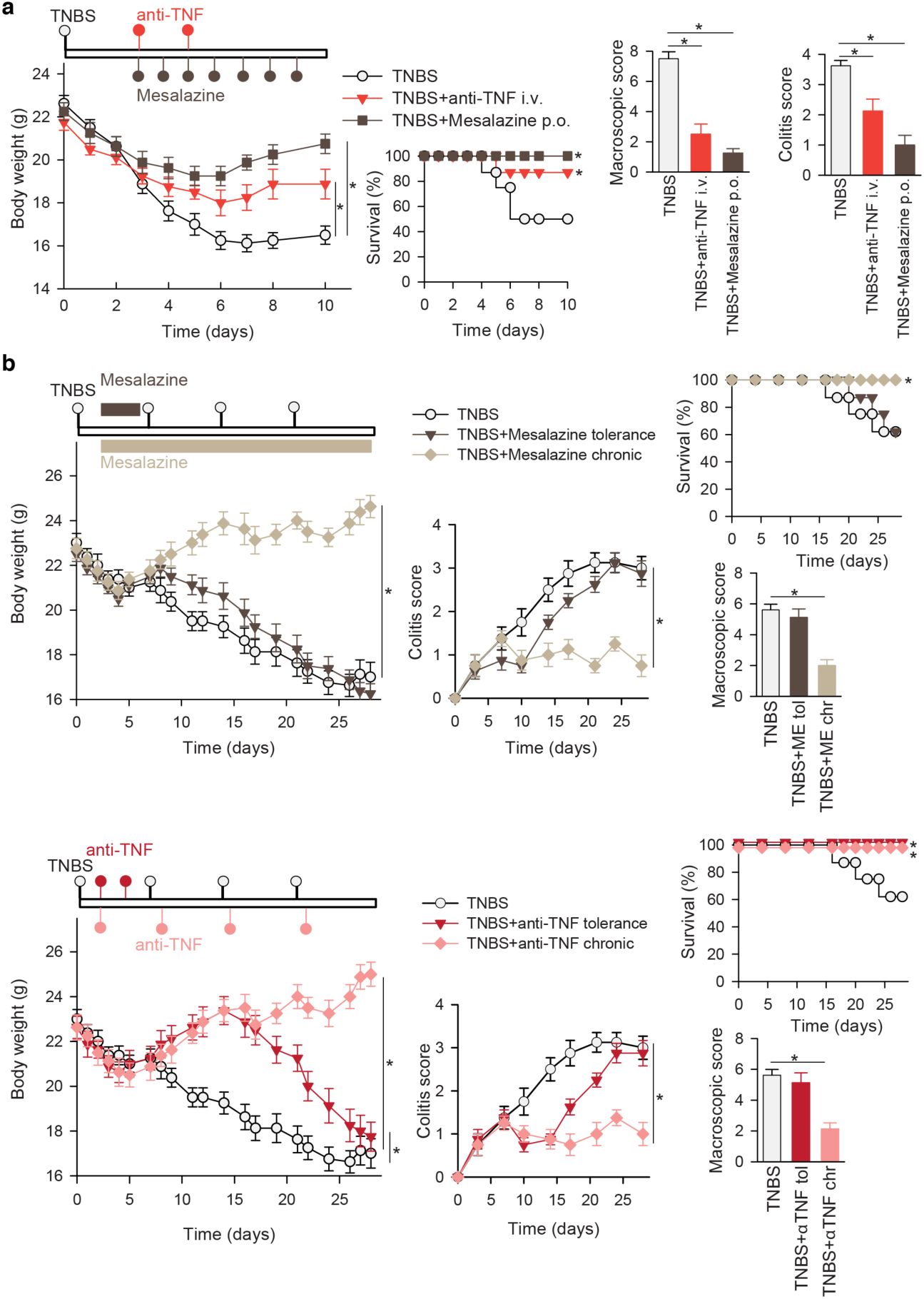
Therapeutic effects of anti-TNFα antibodies and mesalazine on TNBS-induced acute and chronic colitis. **a.** Acute colitis was induced in male Balb/c mice by intrarectal administration of TNBS (3 mg/mouse) in 50% ethanol. Mice were treated in a curative regime with Mesalazine (orally, at 50 mg/kg, twice a day) from days 3 to 6, or with anti-TNFα antibody (intravenously, 5 mg/kg, at days 3 and 5). Disease evolution and severity was monitored by survival and weight loss (expressed in grams). Colitis score (scale 0-4) was determined at day 4 (protective regime) or at day 6 (curative regime). Macroscopic damage score (scale 0-10) was evaluated in colons isolated at day 10 or immediately after death of animal. **b.** Chronic colitis was induced in male Balb/c mice by intrarectal injections of increasing doses of TNBS once a week (0.8 mg/mouse at day 0, 1 mg/mouse at day 7, 1.2 mg/mouse at day 14 and 1.5 mg/mouse at day 21). Mice were treated with mesalazine (orally, 50 mg/kg, twice a day, in 100 μl saline) in a regime of tolerance (from days 3 to 6) or in a chronic treatment (from days 3 to 28), or with anti-mouse TNFα antibody (intravenously, 5 mg/kg) in a treatment of tolerance (at days 3 and 5) or in a chronic treatment (at days 3, 8, 15 and 22). Disease evolution and severity were evaluated by determining body weight loss, survival, colitis score and the macroscopic colonic damage (at day 28 or after death of each animal). n=8 mice/group. *p<0.05 versus untreated TNBS-colitic mice.

**Supplementary Fig. 5.**
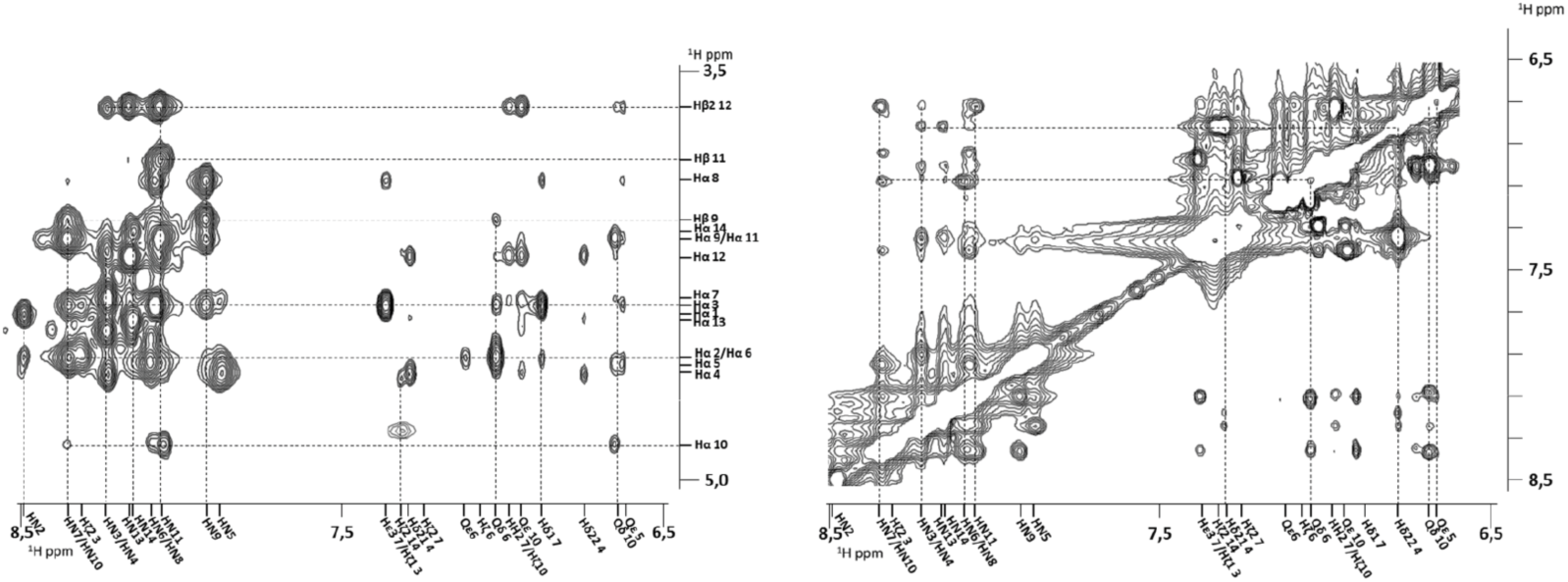
Amide/proton alpha(left) and amide/aromatic (right) regions of the analog **5** (^1^H-Noesy experiment), with peak assignments.

