## Supplementary Material and Methods for "Structure-based design of a Cortistatin analog with improved immunoregulatory activity against inflammatory bowel disease (IBD)"

[d.] *BCN Peptides S.A. Pol.Ind. Els Vinyets-Els Fogars, Sector II. Ctra. Comarcal 244, Km. 22, 08777 Sant Quintí de Mediona, Barcelona08777, Spain.*

[e.] *Institució Catalana de Recerca i Estudis Avançats (ICREA), Passeig Lluis Companys, 23, Barcelona 08010, Spain*

[§] *Present Address: Departament de Ciències Experimentals i de la Salut, Universitat Pompeu Fabra, Dr. Aiguader, 88 08003 Barcelona, Spain*

### Table of Contents

*Methods* ... *3*

1. General syntheses of peptides 1-7 ... 3
2. NMR Methods. ... 3
3. Binding values ... 4
4. Determination of in vitro stability of peptides in human plasma ... 4
5. Inflammatory response *in vitro*. ... 5
6. Immune responses *in vitro*. ... 5
7. Induction of acute chronic ulcerative colitis with dextran sulfate sodium (DSS): 6
8. Induction of acute and chronic colitis with trinitrobenzene sulfonic acid (TNBS) 7
9. Data Analysis ... 8

*Supplementary Figures* ... *9*

Supplementary Figure 1 ... 9

Supplementary Figure 3. ... 10

Supplementary Figure 4. ... 11

Supplementary Figure 5 ... 13

*NMR assignments, spectra and structure calculation statistics.* ... *14*

*Reverse phase – HPLC analysis of compounds* ... *26*

*ESI - MS analysis of compounds* ... *32*

*Bibliography* ... *35*

### Methods

#### 1. General syntheses of peptides 1-7.

The syntheses were performed by SPPS on 0.25 g of 2-Cl-Trt resin (1.60mmol/g) using the Fmoc/tBu strategy. The C-terminal first amino acid (1.0 eq) was coupled in the presence of DIPEA (3.0 eq) in 1.25 ml of DCM as solvent for 45 min and finally end-capped with methanol (0.8 mL·g-1). Then, Fmoc removal was performed by treating the peptidyl resin with 20% piperidine in DMF (1 x 1' and 1 x 5'). The next amino acids (2.5 eq) were coupled using DIPCDI (2.5 eq) and HOBt (2.5 eq), as activating reagents, in DMF for 40-60 min. Kaiser Test was used to check coupling completion. This procedure was repeated for the following Fmoc-protected amino acids and for the last Boc-Pro-OH in compounds **1**, **2** and **4**, and Boc-L-Ala-OH in compound **7**. When Fmoc-L-Msa-OH and Fmoc-L-Dfp-OH were coupled, only 1.5 eq were used. The cleavage of the fully protected linear peptide from the resin was carried out using a cleavage cocktail (DCM:TFE:AcOH, 70:20:10 (v/v)) for 2h. The formation of the disulfide bridge in all the analogues was achieved using iodine (0.73 g, 10 eq) in 3.57 ml of DCM:TFE:AcOH solution at room temperature for 15' and then quenched with an aqueous solution of sodium thiosulphate (1.52 g, 22 eq) in 6,12 ml of water . The aqueous layer was extracted with DCM (3 x 4 mL), the combined organic layer was washed with a mixture of an aqueous citric acid 5% solution/sodium chloride (1:1) and evaporated under reduced pressure. Finally, total deprotection of the side-chains was performed using 6.8 ml of acidic mixture (TFA/DCM/Anisole/$H_2O$, 55:30:10:5 (v/v)) for 4h. Then, the remaining solution was washed with heptane (13 mL) and the aqueous phase was precipitated in $Et_2O$ (-10ºC). The obtained suspension was filtered through a filter plate and the filtrates were discarded. The residue was washed with ether, discarding the filtrates in each washing.

The crude was purified by semipreparative system equipped with a NW50 column filled with silica Kromasil 10 microns. The peptide was suspended in 0.1 N AcOH and DOWEX resin fitted out in 0.1 N AcOH was added. The final acetate compound was recovered by filtration and characterized by mass spectrometry in ESI-MS equipment.

#### 2. NMR Methods.

NMR experiments were recorded at 285 K using a Bruker Avance III 600-MHz spectrometer equipped with a quadruple resonance z-gradient CryoProbe (5 mm

QPQCI). The data was processed TopSpin version 3.1 and chemical shift assignments were generated using Cara[1] and a combination of 2D TOCSY and NOESY homonuclear experiments. Distance restraints derived from the NOESY experiments were used for structure calculation using unambiguously assigned peaks. The structures were calculated with the program CNS 1.2[2]. The protocol consisted of an implicit water simulated-annealing of 120 structures using 8,000 cooling steps followed by an explicit water refinement of the calculated structures using all experimental restraints during 1200 steps (as decribed in[3]).

#### 3. Binding values

All peptides were subjected to anion exchange ($F_3CCOO^- \rightarrow AcO^-$) by using DOWEX resin (DowexMonosphere 550A (OH)). Biological tests were carried out with previously purified peptides containing acetate counter-ion.

Human recombinant CHO-K1 cells were used expressing independently each of the five somatostatin receptors (SSTR1-SSTR5). Cells in HEPES buffer pH 7.4 was incubated with the new analogues of cortistatin in a concentration range of 1 pM to 1 μM for 2-4 h and $^{125}$I-Tyr11 SST14 was used as radioligand and somatostatin-14 as ligand cold. The radioactivity obtained in the absence of SST14 is considered as total binding and that obtained in the presence of 1 μM of SST14 is considered as nonspecific binding. Specific binding was considered as the difference between total and nonspecific, $IC_{50}$ values were determined by a non-linear, least squares regression analysis using MathIQ™ (ID Business Solutions Ltd., UK). The $K_i$ values were calculated using the equation of Cheng and Prusoff[4] using the observed $IC_{50}$ of the tested compound, the concentration of radioligand employed in the assay, and values for the $K_D$ of the ligand (performed at **Eurofins Panlabs, Inc.**). $IC_{50}$, $K_i$, data were performed as duplicates and are presented without Standard Error of the Mean (SEM), therefore, the values presented ($K_i$, $IC_{50}$,) should be interpreted with caution.

#### 4. Determination of in vitro stability of peptides in human plasma

The peptides were dissolved in water at a concentration of 6 mg/mL and warmed up to 37ºC. Human plasma was obtained as a lyophilized solid (K3 EDTA Plasma, BBI solutions, code S112-1), reconstituted with sterile 0.9% sodium chloride solution and stored at -20ºC. Human plasma was thawed and incubated at 37ºC before use.
The peptides were incubated in 90% human plasma at 37ºC for different times and then precipitated with two volume equivalents of methanol. The samples were cooled

in an acetone-carbon dioxide bath for a few seconds and centrifuged for 12 minutes at 4ºC at approximately 10,000 rpm. The supernatant was filtered with a 0.45 µm PVDF filter and analyzed in triplicate by RP-HPLC using an isocratic method (Eluent A=0.1%TFA in water; Eluent B=0.07%TFA in ACN, Column = Kromasil C8, 100 Å, 5 µm, 250 x 4.6mm, Flow = 1 mL/min, Wavelength = 220 nm, Injection volume = 20µL, Tª= 60ºC).The disappearance of the peptide was determined in relation to the area of the initial time for calculating its half-life.

#### 5. Inflammatory response *in vitro*.

RAW 264 cells (obtained from ATTC) were grown at $5x10^5$ cells/ml in complete DMEM medium (DMEM medium supplemented with 100 U/ml penicillin/streptomycin, 2 mM L-glutamine, 50 µM 2-mercaptoethanol and 10% heat-inactivated fetal calf serum) until reaching a confluence of 80% and then incubated in the absence (used as baseline reference) or presence of lipopolysaccharide (LPS, 1 µg / ml, from *E. coli* serotype 055: B5, Sigma). New Cortistatin analogs were added at a concentration of 100 nM at the beginning of culture. After 24 hours of culture, supernatants were harvested and levels of cytokines (TNFα and IL-6) and nitric oxide were determined by ELISA (Preprotech) and the Griess assay, respectively.

#### 6. Immune responses *in vitro*.

Spleen cells were isolated from C57Bl/6 mice (male, 8 weeks of age) by mechanical cell dissociation, filtration using a nylon filter and red cell lysing. The spleen cells were incubated in complete DMEM medium at a density of $10^6$ cells/ml for 2 hours. Nonadherent cells (formed by 80% by T cells) were cultured in complete DMEM and stimulated with anti-CD3 antibody (2 µg/ml) in the presence of different Cortistatin analogs (100 nM). After 48 hours, culture supernatants were collected and the levels of cytokines (IFNγ and IL-2) were determined by ELISA (Preprotech). To determine the effect of different analogues of Cortistatin in proliferation, the cells were cultured for 72 hours, 0.5 µCi (0.0185 MBq)/well of $[^3H]$-thymidine was added for the last 8 hours of culture, harvested the membranes and $[^3H]$-thymidine incorporated was quantified by scintillation counting.

7. Induction of acute chronic ulcerative colitis with dextran sulfate sodium (DSS):

Acute colitis was induced in male C57Bl/6 mice (7-week old, Charles River) by administering 5% dextran sulfate sodium (DSS; molecular weight 20,000 Da; Sigma) from day 0 to day 7 in the drinking water ad libitum. At days 1, 2 and 3, Analog 5 (at 5 nmol/mouse, 0.4 mg/kg, in 100 µl saline) was injected intraperitoneally (i.p.) in the DSS-treated animals. Cortistatin (CST, 5 nmol/mouse) injected i.p. at days 1, 2 and 3 was used as a product of reference.

Recurrent-remitting colitis was induced in male C57Bl/6 mice (7-weeks old, Charles River) by administering 3% DSS (molecular weight 20,000 Da; in the drinking water ad libitum) in two cycles of 7 days (from day 0 to day 7 and from day 15 to day 21 with normal water between cycles). Animals hydrated with normal water were used as naïve controls. Analog 5 (0.4 mg/kg, in 100 µl saline) was injected subcutaneously (s.c.) in the DSS-treated animals following two profiles: during the first cycle of DSS at days 3, 4 and 5; or during the two cycles of DSS at days 3, 4, 5, 16, 17 and 18. A neutralizing anti-mouse TNFα antibody (clone TN3-19.12 from BD Bioscience) injected intravenously (i.v., in 20 µl saline through the tail vein) at days 3, or at days 3 and 16 was used as a treatment of reference.

In both acute and chronic models, colitis severity was assessed daily by scoring (scale 0–4) the clinical disease activity by evaluating stool consistency, presence of faecal blood and weight loss, including a summation of the three components: weight loss (0 = 0%, 0.5 = 1–10%, 1 = 11–15%, 1.5 = 16–20%, 2 = >20%), diarrhea (0 = normal stool, 0.5 = soft stool and minimal wet anal fur/tail, 1 = diarrhea and moderate-to-severe wet anal fur/tail), and frank rectal bleeding (0 = absent, 0.5 = present but minimal, 1 = moderate/severe). Mice were sacrificed on day 8 (acute model) or on day 25 (chronic model), the entire colon was removed from the caecum to the anus, and colon length and weight were measured as indirect inflammation markers. The macroscopic colonic damage score (scale 0-8) was assessed based on the grade of tissue adhesion, presence of ulceration and wall thickness: ulceration (0 = normal appearance, 1 = focal hyperaemia, no ulcers, 2 = ulceration without hyperemia or bowel wall thickening, 3 = ulceration with inflammation at 1 site, 4 = two or more sites of ulceration and inflammation, 5 = major sites of damage extending > 1 cm along length of colon), adhesions (0 = no adhesions, 1 = minor adhesions, colon can be easily separated from the other tissues, 2 = major adhesions), thickness (maximal bowel wall thickness, in mm, measured with a calliper).

#### 8. Induction of acute and chronic colitis with trinitrobenzene sulfonic acid (TNBS)

To induce acute colitis, 2,4,6-trinitrobenzene sulfonic acid (TNBS, 3 mg; Sigma) in 50% ethanol (100 µl) was administered intrarectally in 6-8 weeks-old male BALB/c mice (150 mg/Kg of TNBS). Control mice received 50% ethanol alone. Animals were treated i.p. or s.c. with vehicle (saline) or with Analog 5 (0.4 mg/kg, 9 µg/mouse, in a total volume of 100 µl) 12, 24 and 36 hours (in a protective regime) after instillation of TNBS or 3, 4 and 5 days after TNBS (in a curative regime). In addition, Analog 5 was administered orally in a curative regime (15 mg/kg) twice/day at days 3, 4 and 5. As treatments of reference, anti-TNFα antibody (5 mg/kg, clone TN3-19.12) was injected i.v. at days 3 and 5, and Mesalazine (5-aminosalycilic acid, 50 mg/kg, Sigma) was administered orally twice/day from day 3 to day 10.

To induce chronic colitis, male Balb/c mice (7 weeks-old) were repetitively injected intrarectally with TNBS (0.8 mg in 100 µl 50% ethanol at day 0;1 mg in 100 µl 50% ethanol at day 7; 1.2 mg in 100 µl 50% ethanol at day 14 and 1.5 mg in 100 µl 50% ethanol at day 21). Controls were injected intrarectally with 50% ethanol (100 µl) at days 0, 7, 14 and 21. Analog 5 was injected s.c. (0.4 mg/kg, in total volume of 100 µl saline), following two different regimens of administration: treatment of tolerance at days 3, 4 and 5, and chronic treatment at days 3, 4, 5, 8, 9, 10, 15, 16, 17, 22, 23 and 24. Anti-mouse TNFα antibody and Mesalazine were used as treatments of reference. Mesalazine was administered orally (50 mg/kg, twice a day, in 100 µl saline) in a treatment of tolerance (from days 3 to 6) or in a chronic treatment (from days 3 to 28). Anti-TNFα antibody was injected i.v. (5 mg/kg) in a treatment of tolerance (at days 3 and 5) or in a chronic treatment (at days 3, 8, 15 and 22).

In both TNBS-induced acute and chronic colitis, animals were daily monitored for the appearance of diarrhea, body weight loss, and survival. At different time points, colitis signs were scored in base of stool consistency and rectal bleeding by two blinded observers using the following scale: 0 = normal stool appearance, 1 = slight decrease in stool consistency, 2 = moderate decrease in stool consistency, 3 = moderate decrease in stool consistency and presence of blood in stools, 4 = severe watery diarrhea and moderate/severe blooding in stools. At day 10 (acute model), at day 28 (chronic model) or immediately after death of each animal, colons were collected and evaluated for macroscopic damage (scale 0–10) based on criteria reflecting inflammation (hyperemia, bowel thickening, and extent of ulceration) in a blinded fashion by two researchers: ulceration (0 = normal appearance, 1 = focal hyperemia, no ulcers, 2 = ulceration without hyperemia or bowel wall thickening, 3 = ulceration with inflammation at 1 site, 4 = two or more sites of ulceration and inflammation, 5 = major sites of damage extending > 1 cm along length of colon, 6-10 = when an area of damage extended > 2 cm along length of colon, score is increased by 1 for each additional cm of involvement).

#### 9. Data Analysis

All values are expressed as mean ± SEM of mice/experiment. The differences between groups were analyzed by Mann–Whitney *U* test and, if appropriate, by Kruskal–Wallis ANOVA test. Survival curves were analyzed by the Kaplan–Meier log-rank test. Changes in body weight were compared by use of the Wilcoxon matched-pair signed-rank test.

### Supplementary Figures

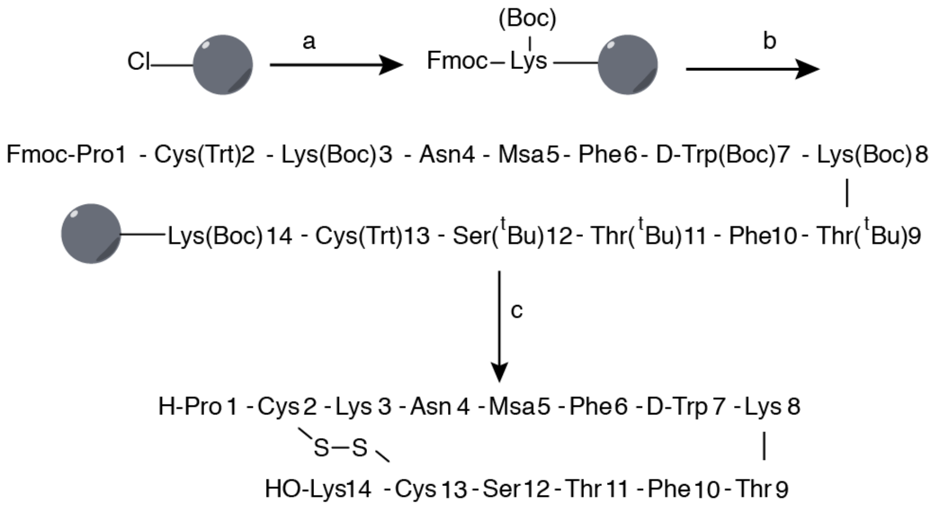


**Supplementary Figure 1.** Synthetic scheme. Synthesis of [L-Msa5_D-Trp7_L- Thr11]-CST14 **(**analog **4).** All peptides have been prepared following a similar approach**.**

a) 1. Fmoc-L-Lys(Boc)-OH (3 equiv) or Fmoc-L-Cys(Trt)-OH (3 equiv), DIEA (3 equiv), 2. MeOH; b) 1. Piperidine 20% DMF, 2. Fmoc-Aaa-OH (1.5–3 equiv, when the Msa amino acid was coupled, only 1.5 equivalents were used), DIPCDI (3 equiv), HOBt (3 equiv), DMF (x12), 3. Piperidine 20% DMF, 4. Fmoc-Pro-OH, DIPCDI, HOBt, DMF

c) 1. $CH_2Cl_2$/TFE/AcOH 2. $I_2$, 3. TFA/$CH_2Cl_2$/anisole/$H_2O$.

Boc=*tert*-butoxycarbonyl, DIEA=diisopropylethylamine, PCDI=diisopropylcarbodiimide, DMF= N,N'-dimethylformamide, Fmoc=N-(9- fluorenylmethoxycarbonyl), HOBT=1-hydroxybenzotriazole, TFA=trifluoroacetic acid, TFE=2,2,2-trifluoroethanol.

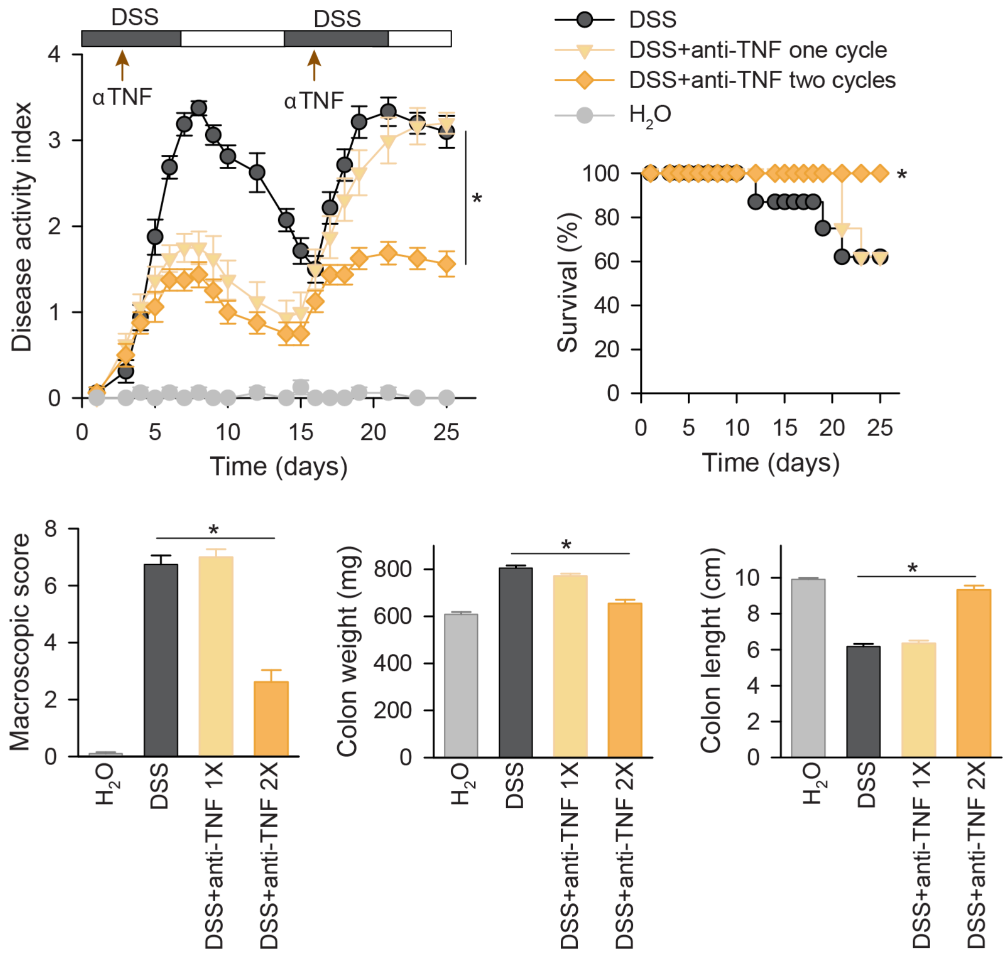


**Supplementary Figure 3. Therapeutic effect of anti-TNFa antibodies in DSS-induced chronic ulcerative colitis.** Mice received 3% DSS with their drinking water in a cyclic manner. Each cycle consisted of 7 days of DSS followed by a 7-day period without DSS supplementation. Anti-mouse TNFα antibody (5 mg/kg) was injected intravenously during the first cycle of DSS at day 3 (1X), or during the two cycles of DSS at days 3 and 16 (2X). Disease activity indexes (scoring body weight loss, stool consistency and presence of fecal blood, scale 0-4) and survival rate were daily evaluated. Signs of colon inflammation were determined at day 25 or immediately after death of each animal by measuring macroscopic damage scores (scale 0-8) and colon length (in cm) and weight (in mg). Mice receiving tap water instead of DSS were used as naïve controls. n=8 mice/group. *p<0.05 versus untreated DSS-colitic mice.

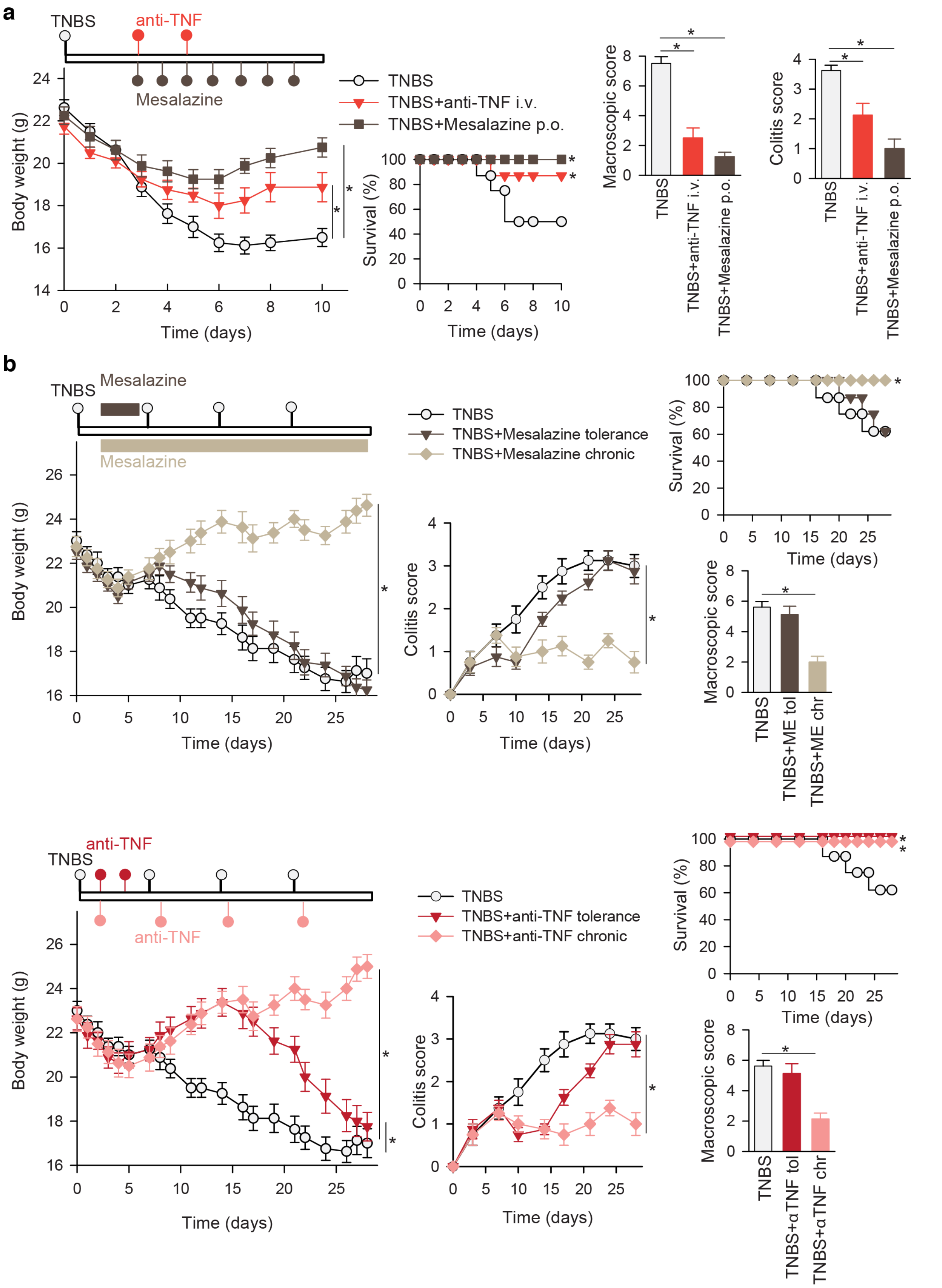


**Supplementary Figure 4. Therapeutic effects of anti-TNFα antibodies and mesalazine on TNBS-induced acute and chronic colitis. A.** Acute colitis was induced in male Balb/c mice by intrarectal administration of TNBS (3 mg/mouse) in 50% ethanol. Mice were treated in a curative regime with Mesalazine (orally, at 50 mg/kg, twice a day) from days 3 to 6, or with anti-TNFα antibody (intravenously, 5 mg/kg, at days 3 and 5). Disease evolution and severity was monitored by survival and weight loss (expressed in grams).

Colitis score (scale 0-4) was determined at day 4 (protective regime) or at day 6 (curative regime). Macroscopic damage score (scale 0-10) was evaluated in colons isolated at day 10 or immediately after death of animal. **B.** Chronic colitis was induced in male Balb/c mice by intrarectal injections of increasing doses of TNBS once a week (0.8 mg/mouse at day 0, 1 mg/mouse at day 7, 1.2 mg/mouse at day 14 and 1.5 mg/mouse at day 21). Mice were treated with mesalazine (orally, 50 mg/kg, twice a day, in 100 μl saline) in a regime of tolerance (from days 3 to 6) or in a chronic treatment (from days 3 to 28), or with anti-mouse TNFα antibody (intravenously, 5 mg/kg) in a treatment of tolerance (at days 3 and 5) or in a chronic treatment (at days 3, 8, 15 and 22). Disease evolution and severity were evaluated by determining body weight loss, survival, colitis score and the macroscopic colonic damage (at day 28 or after death of each animal). n=8 mice/group. *$p<0.05$ versus untreated TNBS-colitic mice.

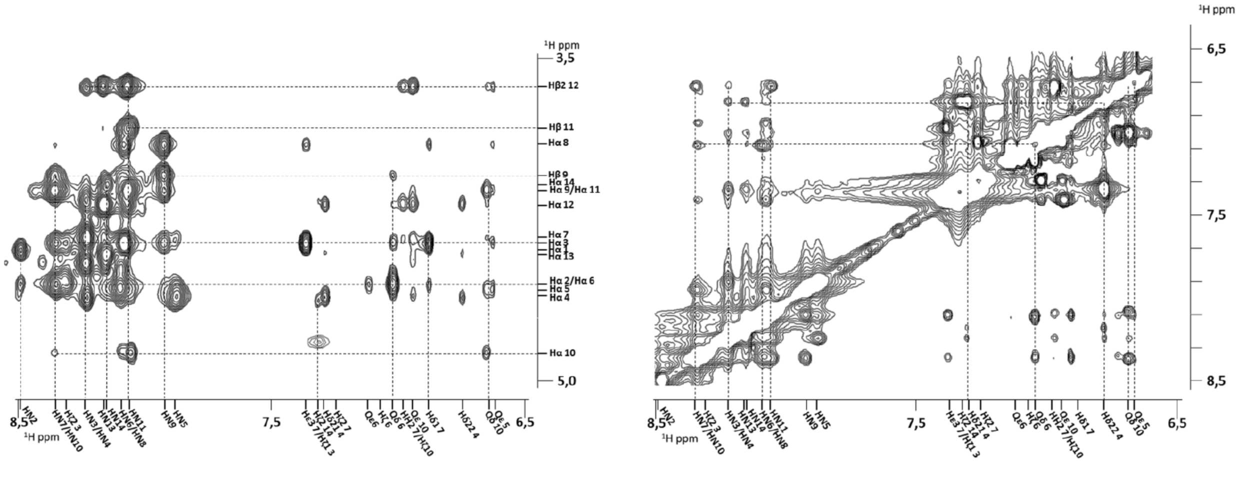


**Supplementary Figure 5.** Amide/proton alpha (left) and amide/aromatic (right) regions of the analog **5** ([1]H-Noesy experiment), with peak assignments.

#### NMR assignments, spectra and structure calculation statistics.

**CST14, 1:** Cortistatin**1** was synthesized following the general procedure from 0.25 g of 2-Cl-Trt resin (1.60 mmol/g) and using Boc-Pro-OH as N-terminal amino acid. ESI-MS: M calcd for 1777.1 g/mol, found (m/z): [M+2H]+/2=889.3, [M+3H]+/3=593.1.

| | HN | Hα | Hβ | Hγ | Hδ | Hε | Hζ | Hη |
|---|---|---|---|---|---|---|---|---|
| **1 Pro** | 7.95 | 4.15 | 2.18 (β2)<br>1.77 (β3) | 1.76 | 3.14 | - | - | - |
| **2 Cys** | 8.67 | 4.43 | 2.90 (β2)<br>2.77 (β3) | - | - | - | - | - |
| **3 Lys** | 8.56 | 4.04 | 1.45 | 1.12 (γ2)<br>1.06 (γ3) | 1.39 | 2.68 | - | - |
| **4 Asn** | 8.11 | 4.38 | 2.34 | - | 7.29 (δ21)<br>6.66 (δ22) | - | - | - |
| **5 Phe** | 8.03 | 4.14 | 2.53 (β2)<br>2.47 (β3) | - | 6.80 | 7.03 | 6.96 | - |
| **6 Phe** | 7.95 | 4.28 | 2.76 (β2)<br>2.69 (β3) | - | 6.89 | 7.05 | - | - |
| **7 Trp** | 7.56 | 4.34 | 2.99 | - | 6.90 | 10.01 (ε1)<br>7.23 (ε3) | 7.17 (ζ2)<br>6.88 (ζ3) | 6.96 |
| **8 Lys** | 7.74 | 3.80 | 1.44 (β2)<br>1.34 (β3) | 0.80 | 1.28 | 2.60 | - | - |
| **9 Thr** | 7.53 | 4.00 | 3.94 | 0.82 | - | - | - | - |
| **10 Phe** | 7.99 | 4.36 | 2.89 (β2)<br>2.75 (β3) | - | 6.95 | 7.05 | - | - |
| **11 Ser** | 8.03 | 4.23 | 3.53 | - | - | - | - | - |
| **12 Ser** | 8.02 | 4.26 | 3.63 (β2)<br>3.60 (β3) | - | - | - | - | - |
| **13 Cys** | 8.23 | 4.47 | 2.94 (β2)<br>2.74 (β3) | - | - | - | - | - |
| **14 Lys** | 7.94 | 3.94 | 1.55 (β2)<br>1.44 (β3) | 1.10 | 1.36 | 2.66 | - | - |

Chemical shifts (ppm) determined using TOCSY and NOESY NMR spectra.

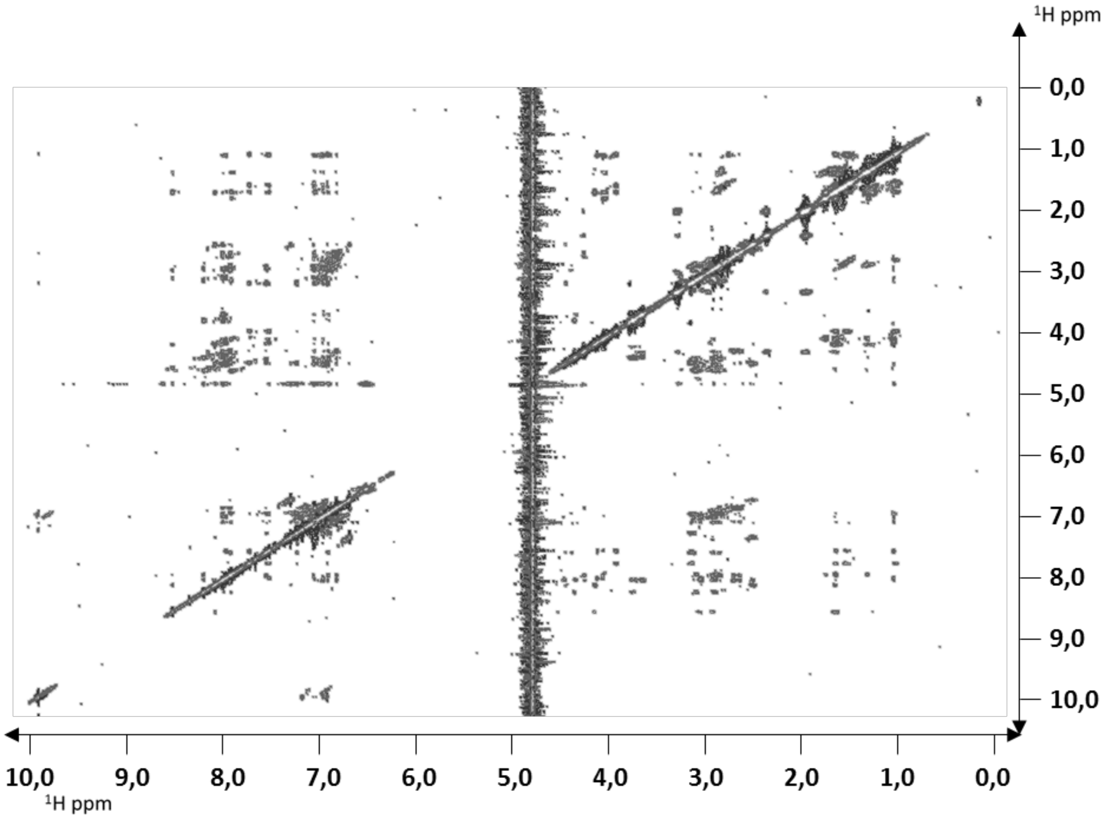


NOESY 350 ms for analog **1**, acquired on a BrukerAvance III 600 MHz a 285 K spectrometer.

**[L-Msa6_D-Trp7_L-Thr11]-CST14, 2:** Cortistatin Analogue **2** was synthesized following the general procedure from 0.25 g of 2-Cl-Trt resin (1.60 mmol/g) and using Boc-Pro-OH as N-terminal amino acid, affording 0.56 g of crude. ESI-MS: M calcd for 1777.1 g/mol, found (m/z): [M+2H]+/2=889.3, [M+3H]+/3=593.1.

| | HN | Hα | Hβ | Hγ | Hδ | Hε | Hζ | Hη |
|---|---|---|---|---|---|---|---|---|
| **1 Pro** | - | 4.16 | 2.21 (β2)<br>2.19 (β3) | 1.81 (γ2)<br>1.77 (γ3) | 3.17 (δ2)<br>3.14 (δ3) | - | - | - |
| **2 Cys** | 8.71 | 4.45 | 2.88 (β2)<br>2.76 (β3) | - | - | - | - | - |
| **3 Lys** | 8.44 | 4.35 | 1.36 | 1.07 (γ2)<br>0.97 (γ3) | 1.23 (δ2)<br>1.13 (δ3) | 2.49 | - | - |
| **4 Asn** | 8.30 | 4.56 | 2.44 (β2)<br>2.35 (β3) | - | 7.33 (δ21)<br>6.75 (δ22) | - | - | - |
| **5 Phe** | 8.03 | 4.14 | 2.53 (β2)<br>2.47 (β3) | - | 6.70 | 6.94 | 6.89 | - |
| **6 Msa** | 8.14 | 4.31 | 2.79 | - | 2.03* | 6.68 | - | 1.74 |
| **7 DTrp** | 8.12 | 4.24 | 2.82 (β2)<br>2.75 (β3) | - | 6.85 | 10.01 (ε1)<br>7.33 (ε3) | 7.25 (ζ2)<br>6.91 (ζ3) | 7.00 |
| **8 Lys** | 8.10 | 3.90 | 1.35 (β2)<br>0.99 (β3) | 0.29 (γ2)<br>0.10 (γ3) | 1.28 | 2.45 (ε2)<br>2.37 (ε3) | - | - |
| **9 Thr** | 7.79 | 4.16 | 3.94 | 0.89 | - | - | - | - |
| **10 Phe** | 8.11 | 4.82 | 2.57 (β2)<br>2.51 (β3) | - | 6.83 | 7.04 | 6.99 | - |
| **11 Thr** | 8.22 | 4.22 | 3.98 | 0.93 | - | - | - | - |
| **12 Ser** | 8.22 | 4.30 | 3.67 | - | - | - | - | - |
| **13 Cys** | 8.26 | 4.38 | 3.66 (β2)<br>2.79 (β3) | - | - | - | - | - |
| **14 Lys** | 7.91 | 3.91 | 1.56 (β2)<br>1.45 (β3) | 1.13 | 1.41 | 2.72 | - | - |

Chemical shifts (ppm) determined using TOCSY and NOESY NMR spectra.

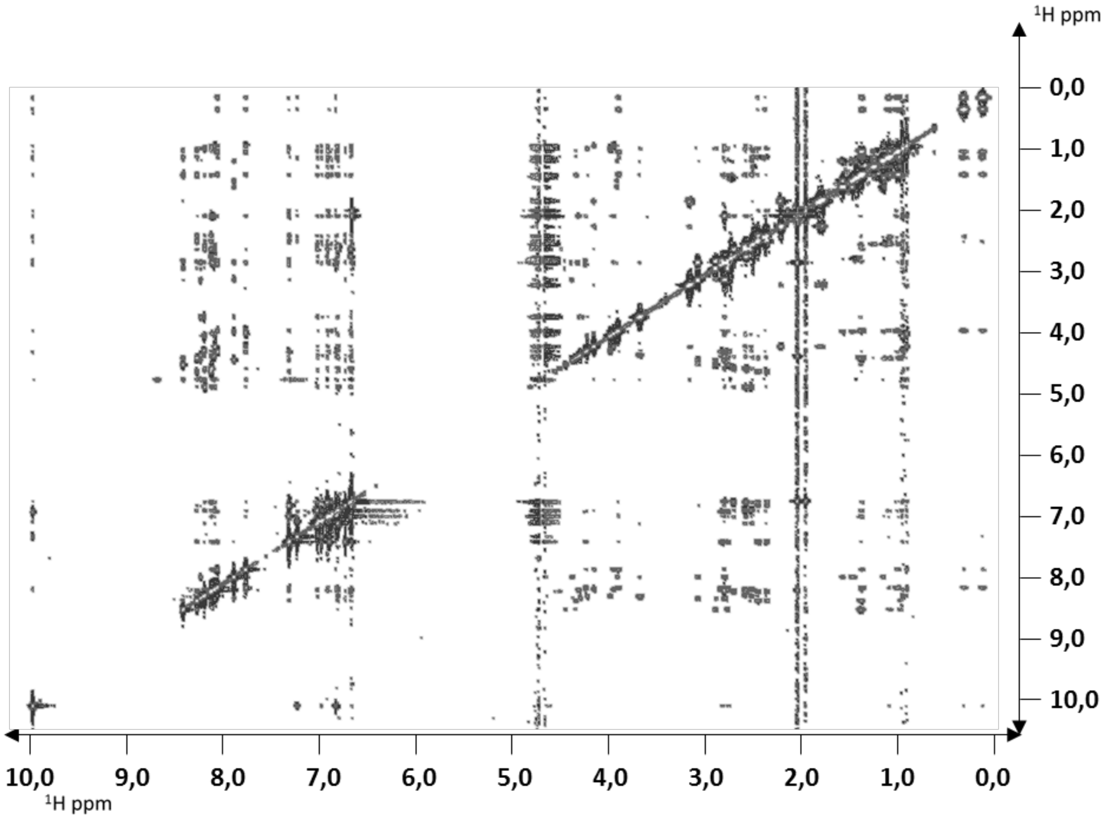


NOESY 350 ms spectrum for analog **2**, acquired on a Bruker Avance III 600 MHz a 285 K spectrometer.

| **DistanceRestricions:** | |
|---|---|
| Intraresidual: 0 | |
| Sequential: 92 | |
| Medium-range (1<dist≤4): 52 | |
| Long-range (dist>4): 56 | |
| Total: 200 | |
| Dihedralanglerestrictions: 24 | |
| Statisticsfor 20 beststructures | |
| **Energies(kcal/mol):** | **RMSD*:** |
| Total energy: 26.09 ± 20.49 | Bonds (Å): 1.000 x10$^{-2}$ ± 4.043x10$^{-4}$ |
| Van der Waals: 106.8 ± 8.184 | Angles (º): 1.806 ± 0.04618 |
| Electrostatic: -495.3 ± 30.44 | Impropers (º): 2.617 ± 0.2759 |
| Bonds: 25.56 ± 2.125 | Dihedrals (º): 43.14 ± 0.1599 |
| Angles: 229.0 ± 12.07 | NOEs: 1.196 x10$^{-2}$ ± 4.760 x10$^{-4}$ |

Structural Statistics for the 20 Lowest Energy Structures of [L-Msa6_D-Trp7_L-Thr11]-CST14 (**2**). *R.m.s deviation between the ensemble of the 20 lowest energy structures and the lowest energy structure.

**Octanoyl-[L-Msa6_D-Trp7_L-Thr11]-CST14 3:** CortistatinAnalogue **3** was synthesized following the general procedure from 0.25 g of 2-Cl-Trt resin (1.60 mmol/g) and using Fmoc-Pro-OH as N-terminal amino acid. Octanoic acid was introduced with 5 eq. of acid, 5 eq. of HOBT and 5 eq. of DIPCDI, yielding 0.50 g of crude. ESI-MS: calcd for 1903.35 g/mol, found (m/z): [M+2H]+/2=952.4, [M+3H]+/2= 635.2.

| | HN | Hα | Hβ | Hγ | Hδ | Hε | Hζ | Hη |
|---|---|---|---|---|---|---|---|---|
| **1 Pro** | - | 4.12 | 2.01 (β2)<br>1.72 (β3) | 1.67 (γ2)<br>1.65 (γ3) | 3.42 (δ2)<br>3.38 (δ3) | - | - | - |
| **2 Cys** | 8.13 | 4.38 | 2.92 (β2)<br>2.73 (β3) | - | - | - | - | - |
| **3 Lys** | 8.28 | 4.34 | 1.39 (β2)<br>1.07 (β3) | 1.34 (γ2)<br>1.20 (γ3) | 0.97 | 2.49 | - | - |
| **4 Asn** | 8.29 | 4.56 | 2.39 | - | 7.33 (δ21)<br>6.76 (δ22) | - | - | - |
| **5 Phe** | 8.13 | 4.49 | 2.71 (β2)<br>2.59 (β3) | - | 6.70 | 6.93 | 6.89 | - |
| **6 Msa** | 8.11 | 4.32 | 2.80 | - | 2.03* | 6.68 | - | 1.95 |
| **7 DTrp** | 8.08 | 4.25 | 2.81 (β2)<br>2.75 (β3) | - | 6.84 | 10.01 (ε1)<br>7.33 (ε3) | 7.25 (ζ2)<br>6.92 (ζ3) | 7.00 |
| **8 Lys** | 8.08 | 3.90 | 1.34 (β2)<br>1.00 (β3) | 0.31 (γ2)<br>0.12 (γ3) | 1.08 | 2.46 (ε2)<br>2.38 (ε3) | - | - |
| **9 Thr** | 7.78 | 4.14 | 3.94 | 0.87 | - | - | - | - |
| **10 Phe** | 8.03 | 4.80 | 2.56 (β2)<br>2.49 (β3) | - | 6.83 | 7.04 | 7.00 | - |
| **11 Thr** | 8.19 | 4.20 | 3.98 | 0.93 | - | - | - | - |
| **12 Ser** | 8.18 | 4.31 | 3.68 | - | - | - | - | - |
| **13 Cys** | 8.15 | 4.29 | 3.08 (β2)<br>2.74 (β3) | - | - | - | - | - |
| **14 Lys** | 7.88 | 3.91 | 1.56 (β2)<br>1.45 (β3) | 1.13 | 1.41 | 2.73 | - | - |

Chemical shifts (ppm) assigned using a pair of TOCSY and NOESY NMR spectra.

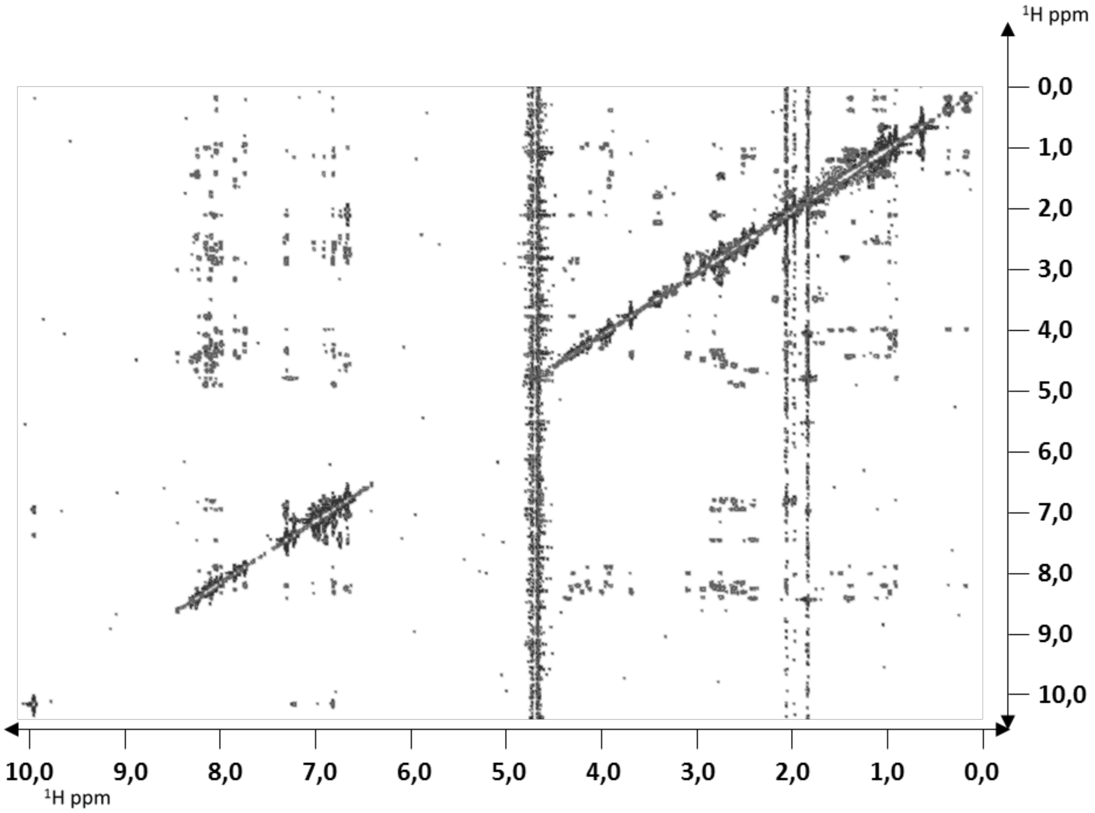


NOESY 350 ms spectrum for analog **3**, acquired on a Bruker Avance III 600 MHz a 285 K spectrometer.

**DistanceRestricions:**

Intraresidual: 0

Sequential: 65

Medium-range (1<dist≤4): 21

Long-range (dist>4): 21

Total: 107

Dihedralanglerestrictions: 0

**Statisticsfor 20 beststructures**

| **Energies (kcal/mol):** | **RMSD*:** |
|---|---|
| Total energy: -273.3 ± 18.53 | Bonds (Å): 3.955 x$10^{-3}$ ± 2,771x$10^{-4}$ |
| Van der Waals: -22.48 ± 10.92 | Angles (º): 1.216 ± 0.03701 |
| Electrostatic: -452.4 ± 33.94 | Impropers (º): 1.430 ± 0.344 |
| Bonds: 4.008 ± 0.5717 | Dihedrals (º): 42.03 ± 1.021 |
| Angles: 103.9 ± 6.411 | NOEs: 6.337 x$10^{-3}$ ± 1,179 x$10^{-3}$ |

Structural Statistics for the 20 Lowest Energy Structures ofoctanoyl-[L-Msa6_D-Trp7_L-Thr11]-CST14 (**3**). *R.m.s deviation between the ensemble of the 20 lowest energy structures and the lowest energy structure.

**[L-Msa5_D-Trp7_L-Thr11]-CST14, 4:**Cortistatin Analogue **4** was synthesized following the general procedure from 0.25 g of 2-Cl-Trt resin (1.60 mmol/g) and using Boc-Pro-OH as N-terminal amino acid, affording 0.53 g of crude. ESI-MS: calcd for 1777.15 g/mol, found (m/z): [M+2H]+/2=889,3 [M+3H]+/2= 593,1.

| | HN | Hα | Hβ | Hγ | Hδ | Hε | Hζ | Hη |
|---|---|---|---|---|---|---|---|---|
| **1 Pro** | 7.74 | 4.16 | 2.21 | 1.81 (γ2)<br>1.78 (γ3) | 3.17 (δ2)<br>3.14 (δ3) | - | - | - |
| **2 Cys** | 8.00 | 4.41 | 2.92 (β2)<br>2.75 (β3) | - | - | - | - | - |
| **3 Lys** | 8.42 | 4.30 | 1.45 | 1.16 (γ2)<br>1.05 (γ3) | 1.35 | 2.64 | - | - |
| **4 Asn** | 8.27 | 4.51 | 2.27 | - | 7.33 (δ21)<br>6.74 (δ22) | - | - | - |
| **5 Msa** | 7.82 | 4.56 | 2.77 (β2)<br>2.68 (β3) | - | 1.93* | 6.65 | - | 1.84 |
| **6 Phe** | 8.16 | 4.46 | 2.79 (β2)<br>2.74 (β3) | - | 7.03 | 7.12 | 7.07 | - |
| **7 DTrp** | 8.42 | 4.29 | 2.84 | - | 6.88 | 10.00 (ε1)<br>7.36 (ε3) | 7.24 (ζ2)<br>6.90 (ζ3) | 6.99 |
| **8 Lys** | 8.11 | 3.87 | 1.37 (β2)<br>0.97 (β3) | 0.25 (γ2)<br>0.06 (γ3) | 1.06 | 2.43 (ε2)<br>2.34 (ε3) | - | - |
| **9 Thr** | 7.97 | 4.05 | 4.03 | 0.90 | - | - | - | - |
| **10 Phe** | 8.46 | 4.76 | 2.64 | - | 6.60 | 6.91 | 6.97 | - |
| **11 Thr** | 8.08 | 4.06 | 3.76 | 0.85 | - | - | - | - |
| **12 Ser** | 8.11 | 4.08 | 3.62 | - | - | - | - | - |
| **13 Cys** | 8.22 | 4.36 | 3.02 (β2)<br>2.74 (β3) | - | - | - | - | - |
| **14 Lys** | 7.86 | 3.91 | 1.55 (β2)<br>1.44 (β3) | 1.12 | 1.40 | 2.71 | - | - |

Chemical shifts (ppm) assigned using a pair of TOCSY and NOESY NMR spectra

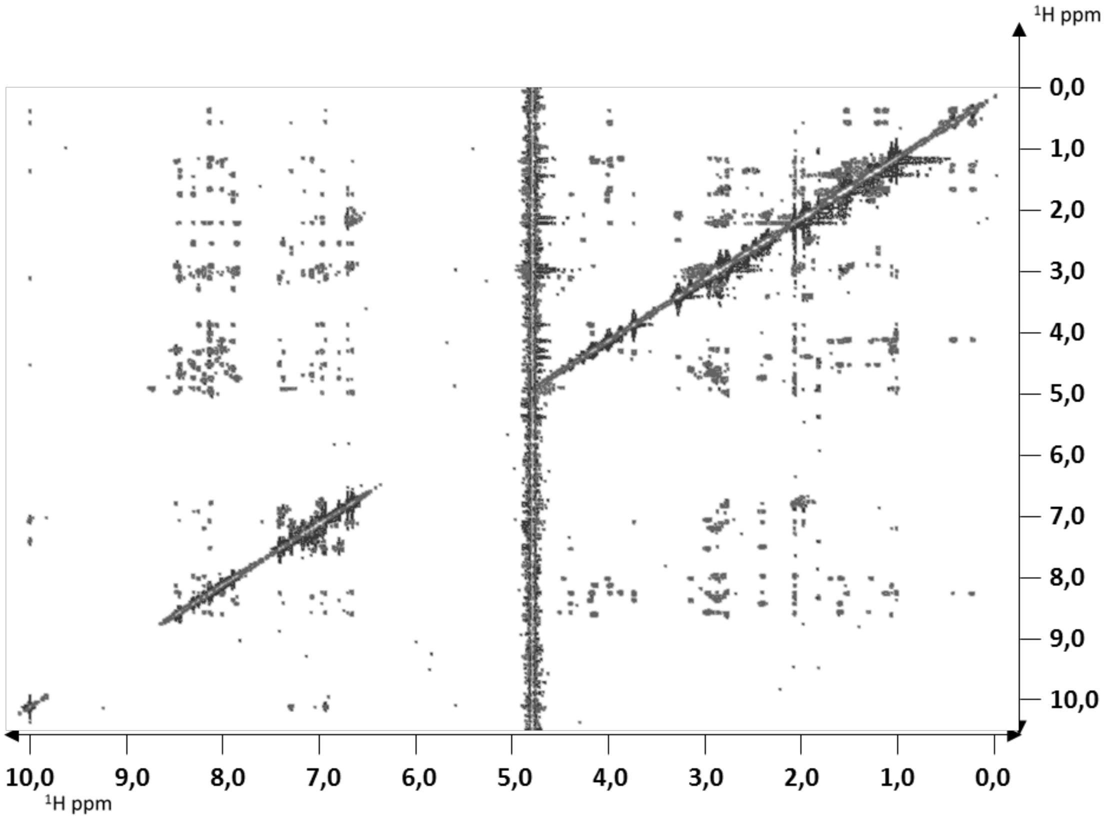


NOESY 350 ms spectrum for analog **4**, acquired on a Bruker Avance III 600 MHz a 285 K spectrometer.

| **DistanceRestricions:** | |
|---|---|
| Intraresidual: 0 | |
| Sequential: 78 | |
| Medium-range (1<dist≤4): 37 | |
| Long-range (dist>4): 50 | |
| Total: 165 | |
| Dihedralanglerestrictions: 20 | |
| **Statisticsfor 20 beststructures** | |
| **Energies (kcal/mol):** | **RMSD*:** |
| Total energy: -77.74 ± 18.24 | Bonds (Å): 6.250 $x10^{-3}$ ± $2.751x10^{-4}$ |
| Van der Waals: 1.073 ± 7.874 | Angles (º): 1.347 ± 0.03784 |
| Electrostatic: -404.2 ± 17.43 | Impropers (º): 4.179 ± 0.2102 |
| Bonds: 10.06 ± 0.8831 | Dihedrals (º): 43.40 ± 0.5269 |
| Angles: 128.3 ± 7.282 | NOEs: 7.625 $x10^{-3}$ ± 5.385 $x10^{-4}$ |

Structural Statistics for the 20 Lowest Energy Structures of [L-Msa5_D-Trp7_L-Thr11]-CST14 (**4**). *R.m.s deviation between the ensemble of the 20 lowest energy structures and the lowest energy structure.

**Octanoyl-[L-Msa5_D-Trp7_L-Thr11]-CST14, 5:** Cortistatin Analogue **5** was synthesized following the general procedure from 0.25 g of 2-Cl-Trt resin (1.60 mmol/g) and using Fmoc-Pro-OH as N-terminal amino acid, and octanoic acid was introduced with 5 eq. of acid, 5 eq. of HOBT and 5 eq. of DIPCDI, yielding 0.55 g of crude. ESI-MS: calcd for 1903.35 g/mol, found (m/z): [M+2H]+/2=952.4, [M+3H]+/2= 635.2.

| | HN | Hα | Hβ | Hγ | Hδ | Hε | Hζ | Hη |
|---|---|---|---|---|---|---|---|---|
| **1 Pro** | - | 4.12 | 2.13 (β2)<br>1.72 (β3) | 1.66 | 3.38 | - | - | - |
| **2 Cys** | 8.15 | 4.37 | 2.88 (β2)<br>2.78 (β3) | - | - | - | - | - |
| **3 Lys** | 8.23 | 4.26 | 1.44 | 1.14 (γ2)<br>1.04 (γ3) | 1.30 | 2.61 | 7.31 | - |
| **4 Asn** | 8.23 | 4.56 | 2.31 | - | 7.28 (δ21)<br>6.73 (δ22) | - | - | - |
| **5 Msa** | 7.88 | 4.55 | 2.72 | - | 1.90* | 6.61 | - | 1.82 |
| **6 Phe** | 8.16 | 4.46 | 2.75 | - | 7.01 | 7.10 | 7.06 | - |
| **7 DTrp** | 8.35 | 4.27 | 2.83 | - | 6.86 | 9.97 (ε1)<br>7.35 (ε3) | 7.23 (ζ2)<br>6.91 (ζ3) | 6.98 |
| **8 Lys** | 8.08 | 3.87 | 1.35 (β2)<br>0.94 (β3) | 0.28 (γ2)<br>0.10 (γ3) | 1.07 | 2.45 (ε2)<br>2.35 (ε3) | 7.23 | - |
| **9 Thr** | 7.92 | 4.06 | 4.00 | 0.89 | - | - | - | - |
| **10 Phe** | 8.36 | 4.29 | 2.62 | - | 6.63 | 6.93 | 6.67 | - |
| **11 Thr** | 8.05 | 4.06 | 3.80 | 0.85 | - | - | - | - |
| **12 Ser** | 8.07 | 4.11 | 3.63 | - | - | - | - | - |
| **13 Cys** | 8.16 | 4.33 | 3.01 (β2)<br>2.72 (β3) | - | - | - | - | - |
| **14 Lys** | 8.15 | 4.03 | 1.49 | 1.41 (γ2)<br>1.16 (γ3) | 1.63 | 2.72 | - | - |

Chemical shifts (ppm) assigned using a pair of TOCSY and NOESY NMR spectra.

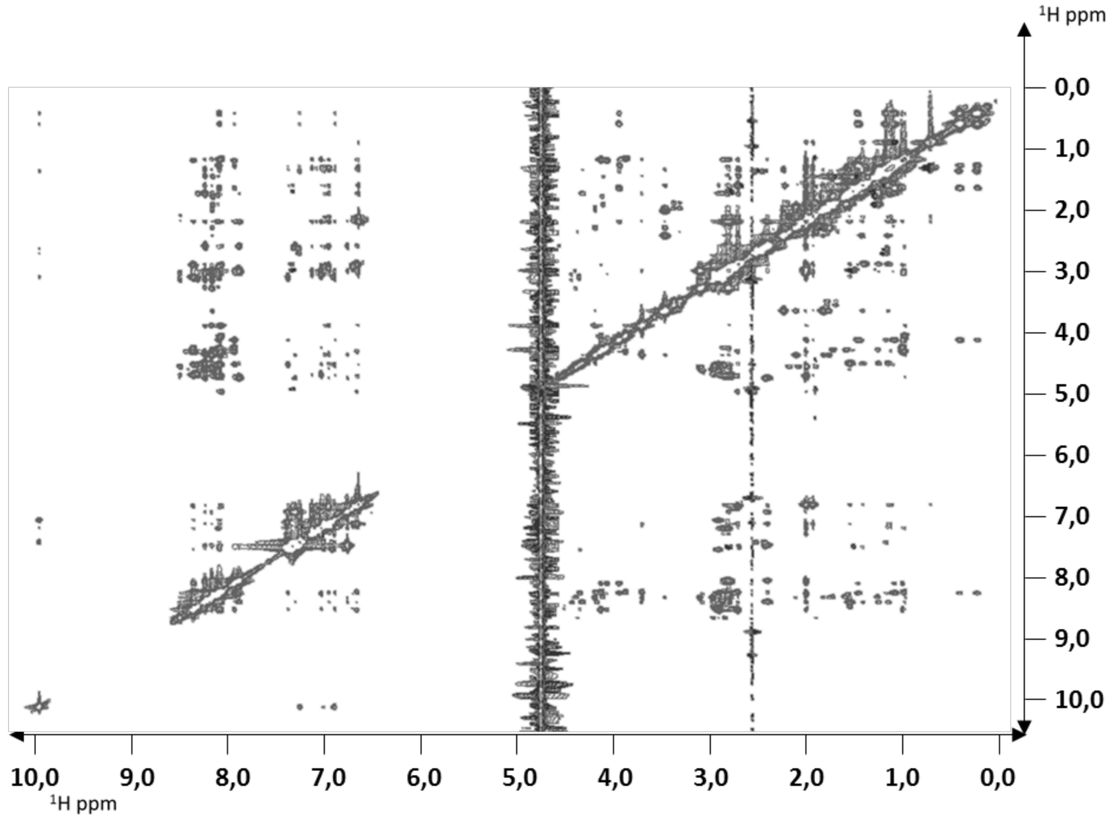


NOESY 350 ms spectrum for analog **5**, acquired on a Bruker Avance III 600 MHz a 285 K spectrometer.

| **DistanceRestricions:** | |
|---|---|
| Intraresidual: 0 | |
| Sequential: 93 | |
| Medium-range (1<dist≤4): 31 | |
| Long-range (dist>4): 49 | |
| Total: 173 | |
| Dihedralanglerestrictions: 24 | |
| **Statisticsfor 20 beststructures** | |
| **Energies (kcal/mol):** | **RMSD*:** |
| Total energy: 12.64 ± 24.44 | Bonds (Å): 8.722 x$10^{-3}$ ± 4.603x$10^{-4}$ |
| Van der Waals: 73.87 ± 12.69 | Angles (º): 1.543 ± 0.04689 |
| Electrostatic: -424.0 ± 28.05 | Impropers (º): 3.284 ± 0.2722 |
| Bonds: 19.45 ± 2.025 | Dihedrals (º): 40.96 ± 0.2467 |
| Angles: 167.3 ± 10.22 | NOEs: 1.123 x$10^{-2}$ ± 7.214 x$10^{-4}$ |

Structural Statistics for the 20 Lowest Energy Structures ofoctanoyl-[L-Msa5_D-Trp7_L-Thr11]-CST14 (**5**). *R.m.s deviation between the ensemble of the 20 lowest energy structures and the lowest energy structure.

**Octanoyl-[L-Msa5_D-Trp7_L-Thr11]-CST13, 6:** Cortistatin Analogue **6** was synthesized following the general procedure from 0.25 g of 2-Cl-Trt resin (1.60 mmol/g) and using Fmoc-Pro-OH as N-terminal amino acid, and octanoic acid was introduced with 5 eq. of acid, 5 eq. of HOBT and 5 eq. of DIPCDI, affording 0.53 g of crude. ESI-MS: M calcd for 1775.18 g/mol, found (m/z): [M+2H]+/2=888.6, [M+3H]+/3= 592.7.

| | HN | Hα | Hβ | Hγ | Hδ | Hε | Hζ | Hη |
|---|---|---|---|---|---|---|---|---|
| **1 Pro** | - | 4.11 | 2.01 | 1.73 (γ2)<br>1.66 (γ3) | 3.40 | - | - | - |
| **2 Cys** | 8.40 | 4.29 | 2.85 | - | - | - | - | - |
| **3 Lys** | 8.25 | 4.30 | 1.43 | 1.15 (γ2)<br>1.02 (γ3) | 1.29 (δ2)<br>1.22 (δ3) | 2.58 | - | - |
| **4 Asn** | 8.24 | 4.55 | 2.30 | - | 7.27 (δ21)<br>6.73 (δ22) | - | - | - |
| **5 Msa** | 8.18 | 4.47 | 2.79 (β2)<br>2.69 (β3) | - | 1.91* | 6.62 | - | 1.80 |
| **6 Phe** | 8.18 | 4.47 | 2.79 (β2)<br>2.73 (β3) | - | 7.03 | 7.11 | 7.07 | - |
| **7 DTrp** | 7.86 | 4.16 | 2.84 | - | 6.88 | 9.99 (ε1)<br>7.36 (ε3) | 7.24 (ζ2)<br>6.90 (ζ3) | 6.99 |
| **8 Lys** | 7.85 | 2.91 | 1.25 | 1.03 | 1.34 | 1.98 | - | - |
| **9 Thr** | 7.93 | 4.09 | 4.00 | 0.89 | - | - | - | - |
| **10 Phe** | 8.33 | 4.09 | 2.55 | - | 6.59 | 6.93 | 6.90 | - |
| **11 Thr** | 8.09 | 4.08 | 3.88 | 0.97 | - | - | - | - |
| **12 Ser** | 8.06 | 4.08 | 3.65 | - | - | - | - | - |
| **13 Cys** | 8.12 | 4.34 | 2.91 (β2)<br>2.73 (β3) | - | - | - | - | - |

Chemical shifts (ppm) assigned using a pair of TOCSY and NOESY NMR spectra.

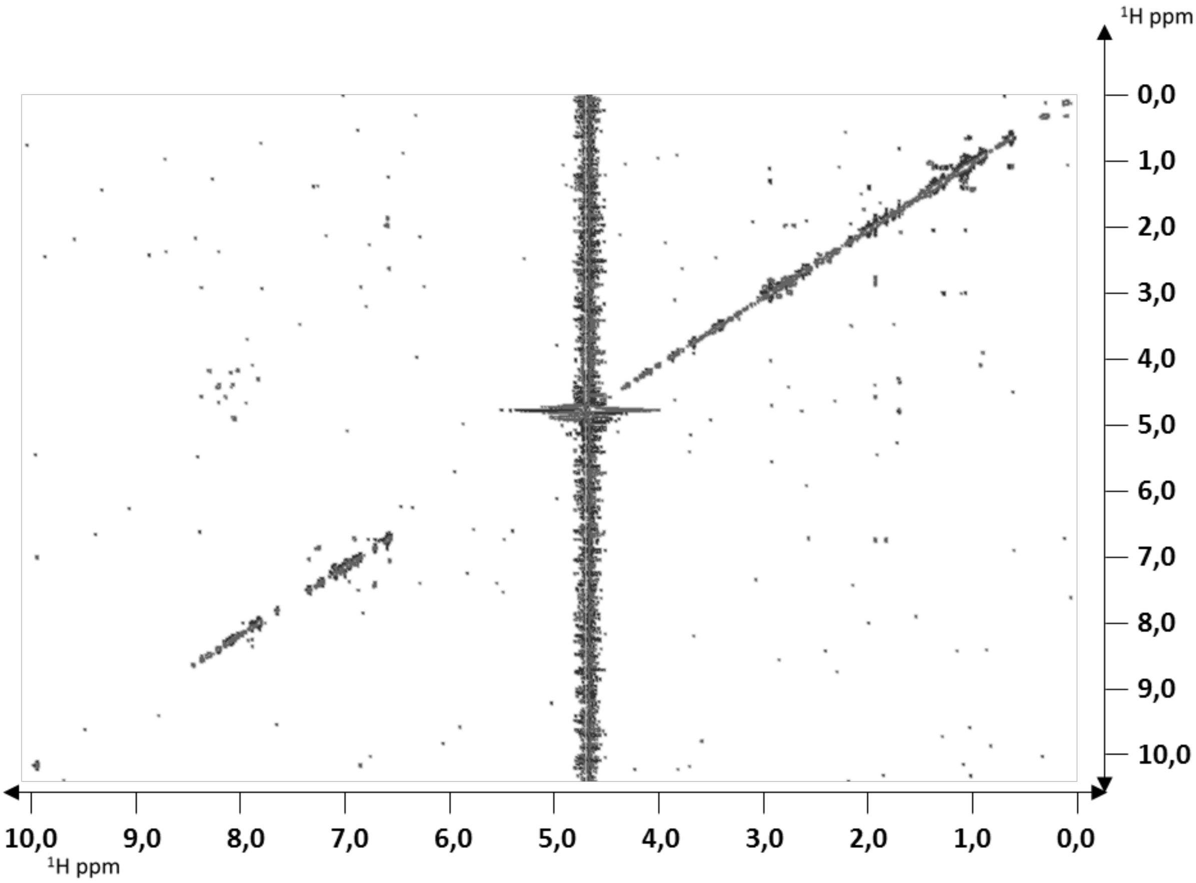


NOESY 350 ms spectrum for analog **6**, acquired on a Bruker Avance III 600 MHz a 285 K spectrometer.

### Reverse phase – HPLC analysis of compounds

All analogues were dissolved in water 1 mg/mL and analysed by RP-HPLC.

RP-HPLC conditions:
Gradient: 5-85%B in 20 min, A: 0.01%TFA in H2O, B: 0.07% TFA in CAN
Column: Kromasil C8, 250 x 4.5, 5um
Wavelength: 220 nm; Flow: 1 mL/min, Tª: 25ºC
Equipment LC20 Shimadzu

#### Analog 1 - RP - HPLC

Eluent A: H2O 0.1% TFA (RD_0605/5)
Eluent B: ACN 0.07% TFA (RD_0606/2)
Columna: Kromasil C8, 250x4.6, 5um, E159909
l:220nm; fluxe:1mL/min; Tª:25ºC
HPLC15

Chromatogram
A61 S1001 @D:\LabSolutions\Data\R+D\Analegs-SOM\SOM-610.lcd

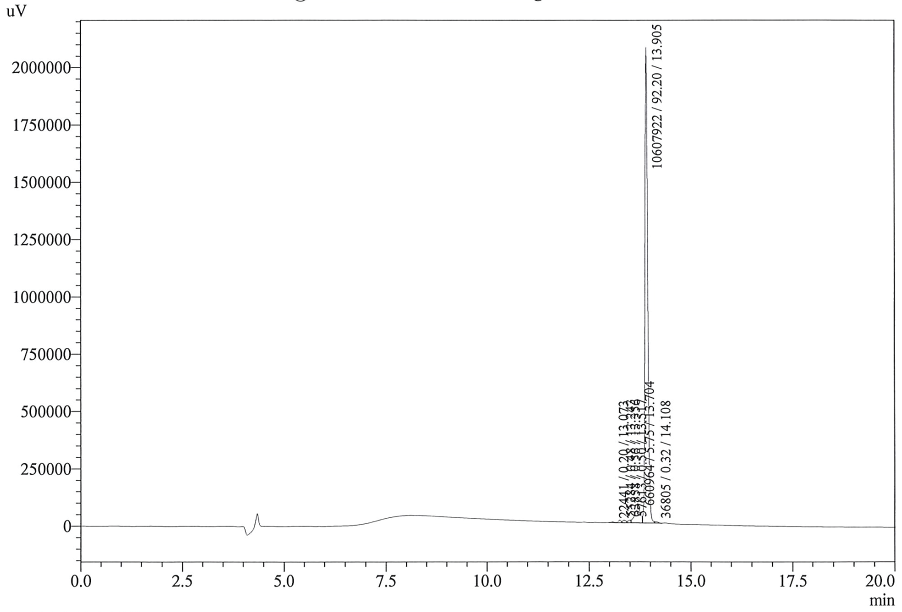


1 Det.A Ch1 / 220nm

PeakTable @D:\LabSolutions\Data\R+D\Analegs-SOM\SOM-610.lcd

Detector A Ch1 220nm

| Peak# | Ret. Time | Area | Area % |
|---|---|---|---|
| 1 | 13.07 | 22441 | 0.20 |
| 2 | 13.24 | 55284 | 0.48 |
| 3 | 13.36 | 63858 | 0.56 |
| 4 | 13.52 | 57613 | 0.50 |
| 5 | 13.70 | 660964 | 5.75 |
| 6 | 13.91 | 10607922 | 92.20 |
| 7 | 14.11 | 36805 | 0.32 |
| Total | | 11504887 | 100.00 |

### Analog 2 - RP - HPLC

Eluent A: H2O 0.1% TFA (RD_0605/5)
Eluent B: ACN 0.07% TFA (RD_0606/2)
Columna: Kromasil C8, 250x4.6, 5um, E159909
l:220nm; fluxe:1mL/min; Tª:25ºC
HPLC15

Chromatogram
A63 15X1 @D:\LabSolutions\Data\R+D\Analegs-SOM\SOM-612.lcd

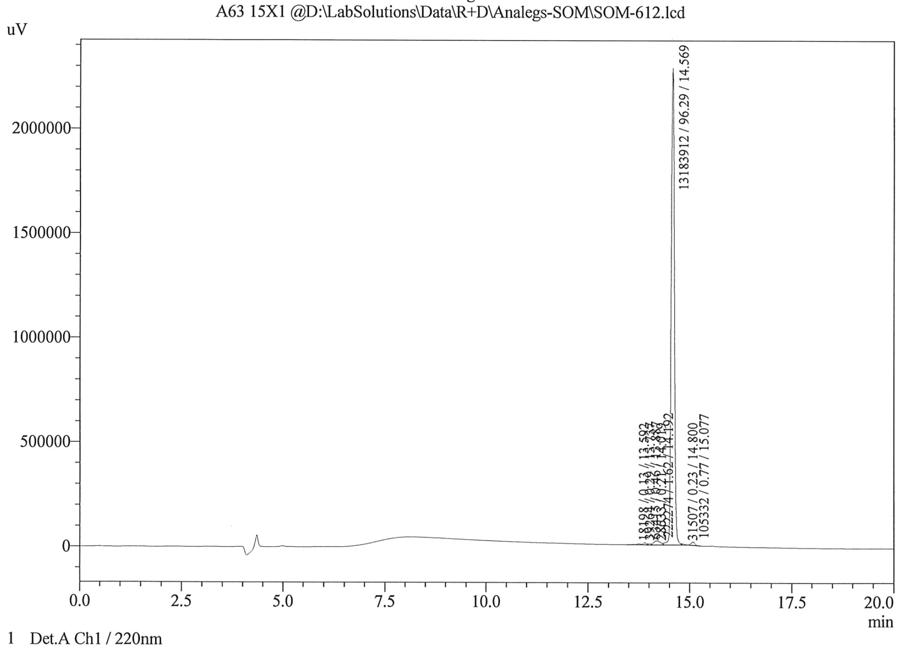


1 Det.A Ch1 / 220nm

PeakTable @D:\LabSolutions\Data\R+D\Analegs-SOM\SOM-612.lcd

Detector A Ch1 220nm

| Peak# | Ret. Time | Area | Area % |
|---|---|---|---|
| 1 | 13.59 | 18198 | 0.13 |
| 2 | 13.74 | 39264 | 0.29 |
| 3 | 13.89 | 63415 | 0.46 |
| 4 | 14.02 | 28633 | 0.21 |
| 5 | 14.19 | 222274 | 1.62 |
| 6 | 14.57 | 13183912 | 96.29 |
| 7 | 14.80 | 31507 | 0.23 |
| 8 | 15.08 | 105332 | 0.77 |
| Total | | 13692535 | 100.00 |

### Analog 3 - RP - HPLC

Eluent A: H2O 0.1% TFA (RD_0605/5)
Eluent B: ACN 0.07% TFA (RD_0606/2)
Columna: Kromasil C8, 250x4.6, 5um, E159909
l:220nm; fluxe:1mL/min; Tª:25°C
HPLC15

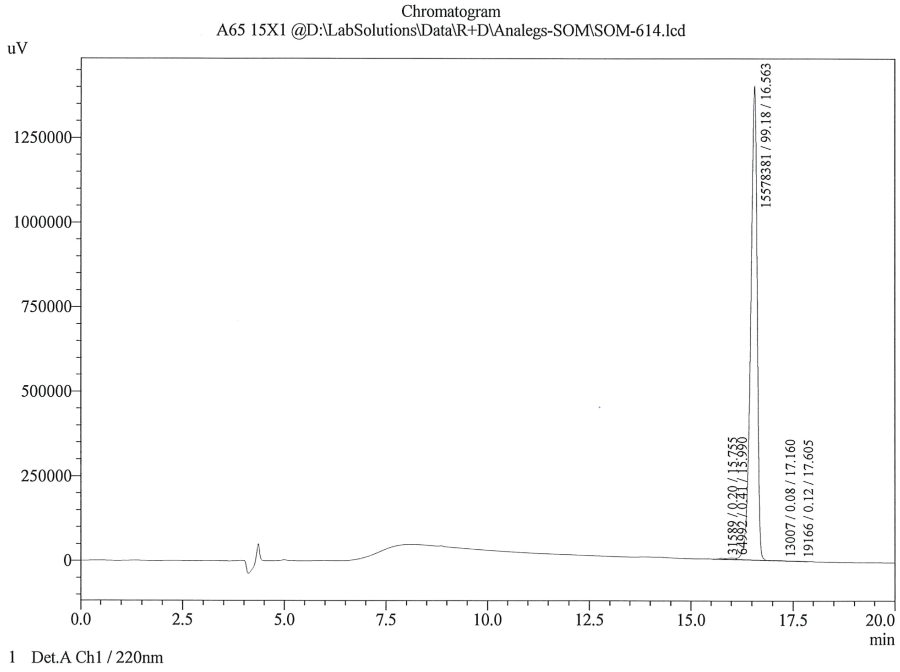


PeakTable @D:\LabSolutions\Data\R+D\Analegs-SOM\SOM-614.lcd

Detector A Ch1 220nm

| Peak# | Ret. Time | Area | Area % |
|---|---|---|---|
| 1 | 15.75 | 31589 | 0.20 |
| 2 | 15.99 | 64992 | 0.41 |
| 3 | 16.56 | 15578381 | 99.18 |
| 4 | 17.16 | 13007 | 0.08 |
| 5 | 17.60 | 19166 | 0.12 |
| Total | | 15707134 | 100.00 |

### Analog 4 - RP - HPLC

Eluent A: H2O 0.1% TFA (RD_0605/5)
Eluent B: ACN 0.07% TFA (RD_0606/2)
Columna: Kromasil C8, 250x4.6, 5um, E159909
l:220nm; fluxe:1mL/min; Tª:25°C
HPLC15

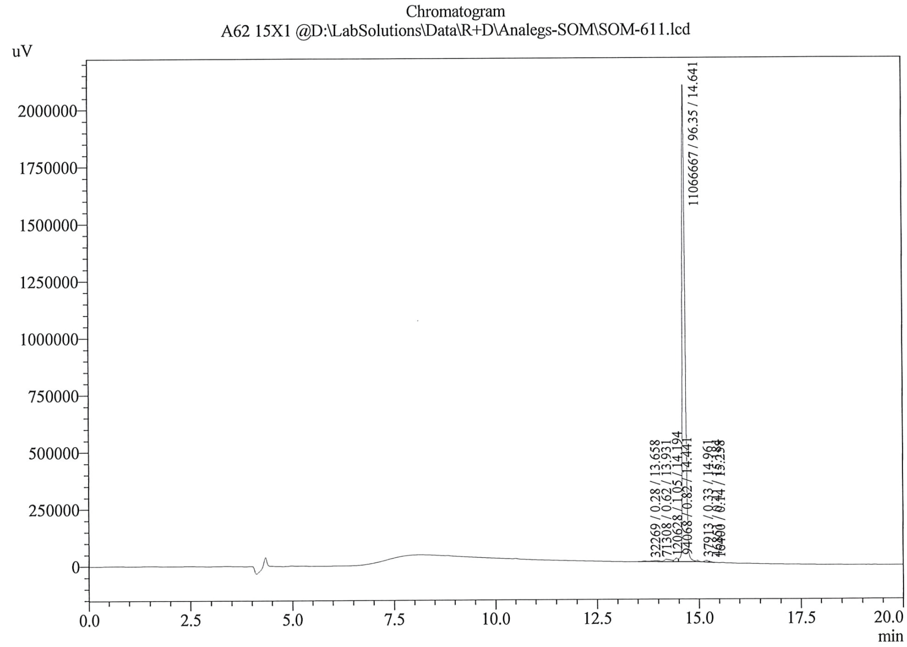


1 Det.A Ch1 / 220nm

PeakTable @D:\LabSolutions\Data\R+D\Analegs-SOM\SOM-611.lcd

Detector A Ch1 220nm

| Peak# | Ret. Time | Area | Area % |
|---|---|---|---|
| 1 | 13.66 | 32269 | 0.28 |
| 2 | 13.93 | 71308 | 0.62 |
| 3 | 14.19 | 120628 | 1.05 |
| 4 | 14.44 | 94068 | 0.82 |
| 5 | 14.64 | 11066667 | 96.35 |
| 6 | 14.96 | 37913 | 0.33 |
| 7 | 15.18 | 46851 | 0.41 |
| 8 | 15.26 | 16400 | 0.14 |
| Total | | 11486103 | 100.00 |

### Analog 5 - RP - HPLC

Eluent A: H2O 0.1% TFA (RD_0605/5)
Eluent B: ACN 0.07% TFA (RD_0606/2)
Columna: Kromasil C8, 250x4.6, 5um, E159909
l:220nm; fluxe:1mL/min; Tª:25ºC
HPLC15

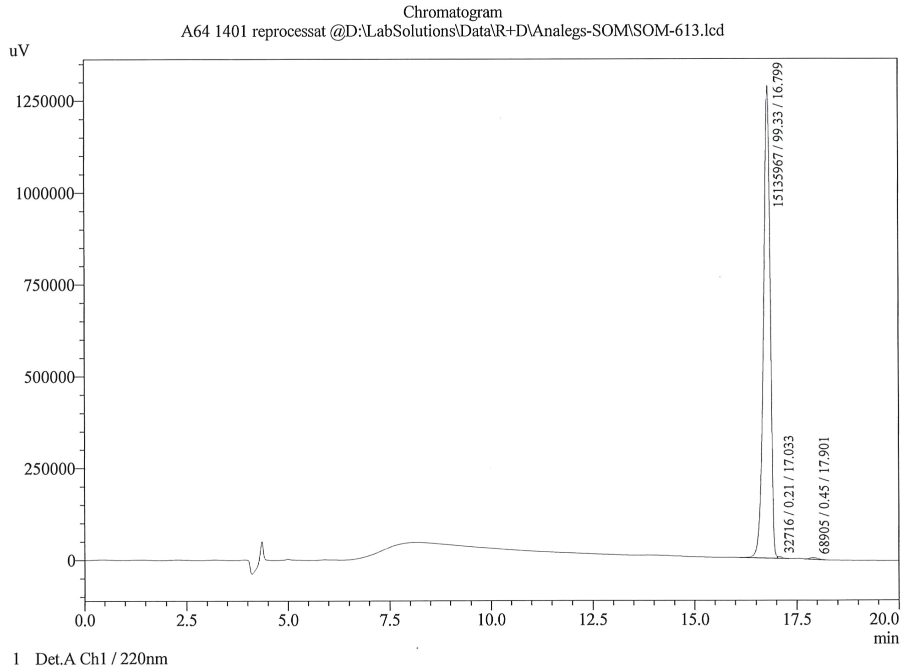


PeakTable @D:\LabSolutions\Data\R+D\Analegs-SOM\SOM-613.lcd

Detector A Ch1 220nm

| Peak# | Ret. Time | Area | Area % |
|---|---|---|---|
| 1 | 16.80 | 15135967 | 99.33 |
| 2 | 17.03 | 32716 | 0.21 |
| 3 | 17.90 | 68905 | 0.45 |
| Total | | 15237588 | 100.00 |

### Analog 6 - RP - HPLC

Eluent A: H2O 0.1% TFA (RD_0605/5)
Eluent B: ACN 0.07% TFA (RD_0606/2)
Columna: Kromasil C8, 250x4.6, 5um, E159909
l:220nm; fluxe:1mL/min; Tª:25°C
HPLC15

Chromatogram
A69 14X1 @D:\LabSolutions\Data\R+D\Analegs-SOM\SOM-615.lcd

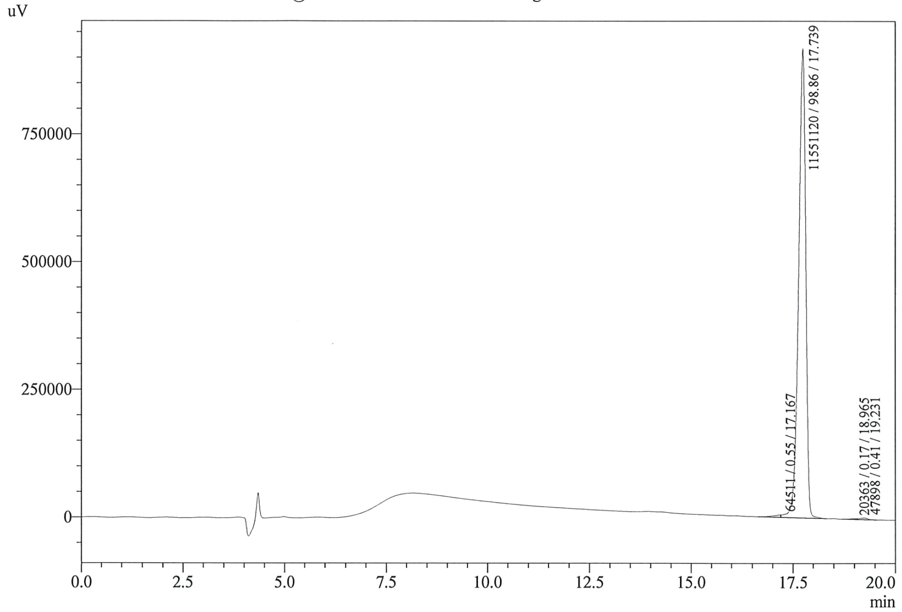


1 Det.A Ch1 / 220nm

PeakTable @D:\LabSolutions\Data\R+D\Analegs-SOM\SOM-615.lcd

Detector A Ch1 220nm

| Peak# | Ret. Time | Area | Area % |
|---|---|---|---|
| 1 | 17.17 | 64511 | 0.55 |
| 2 | 17.74 | 11551120 | 98.86 |
| 3 | 18.97 | 20363 | 0.17 |
| 4 | 19.23 | 47898 | 0.41 |
| Total | | 11683893 | 100.00 |

### ESI - MS analysis of compounds

All analogues were dissolved in water 1 mg/mL and analysed by Electrospray Ionisation Mass Spectrometry..

ESI-MS conditions:
A: 0.01%TFA in H2O, B: 0.07% TFA in ACN
Equipment LCMS8040 Shimadzu

#### Analog 1 - ESI-MS

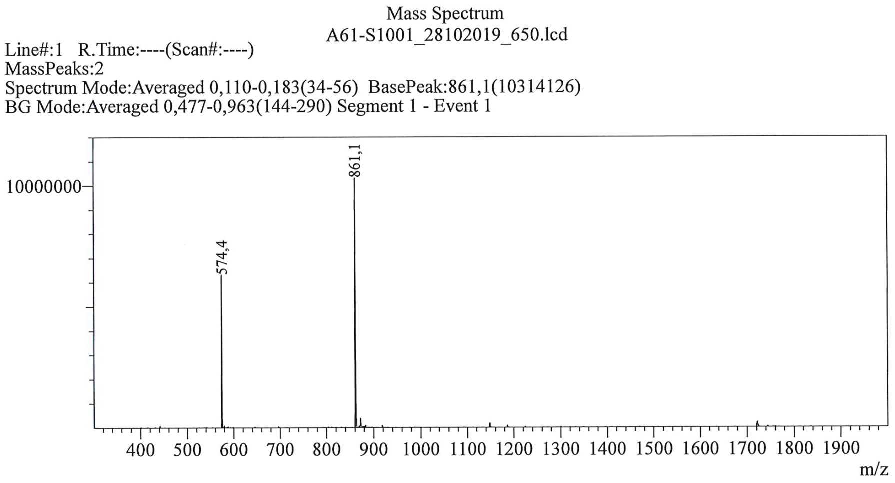


#### Analog 2 - ESI-MS

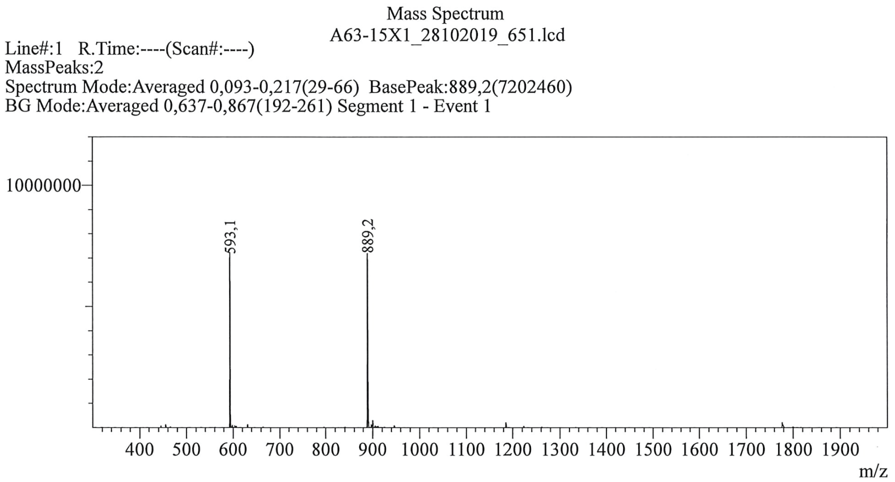

### Analog 3 - ESI-MS

Mass Spectrum
A65-15X1_28102019_653.lcd
Line#:1 R.Time:----(Scan#:----)
MassPeaks:2
Spectrum Mode:Averaged 0,100-0,207(31-63) BasePeak:952,3(8885292)
BG Mode:Averaged 0,557-0,827(168-249) Segment 1 - Event 1

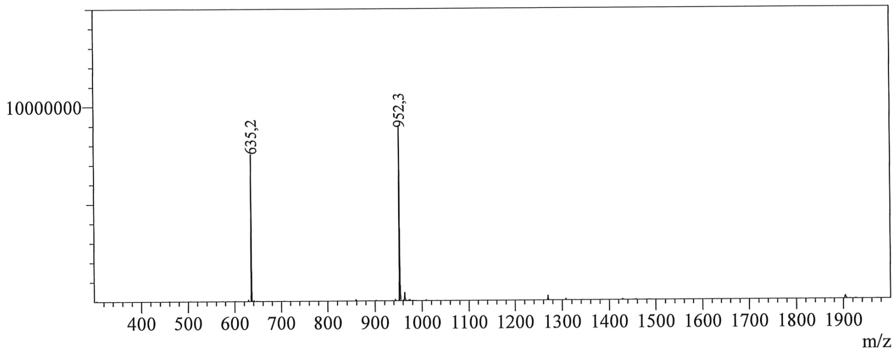


### Analog 4 - ESI-MS

Mass Spectrum
A62-15X1_28102019_649.lcd
Line#:1 R.Time:----(Scan#:----)
MassPeaks:2
Spectrum Mode:Averaged 0,093-0,207(29-63) BasePeak:889,1(7063200)
BG Mode:Averaged 0,427-0,913(129-275) Segment 1 - Event 1

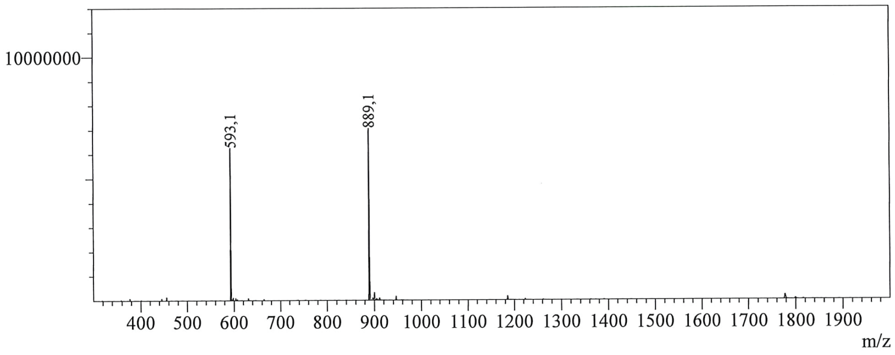

### Analog 5 - ESI-MS

Mass Spectrum
A64-1401_28102019_654.lcd
Line#:1 R.Time:----(Scan#:----)
MassPeaks:2
Spectrum Mode:Averaged 0,097-0,213(30-65) BasePeak:952,2(9751288)
BG Mode:Averaged 0,430-0,843(130-254) Segment 1 - Event 1

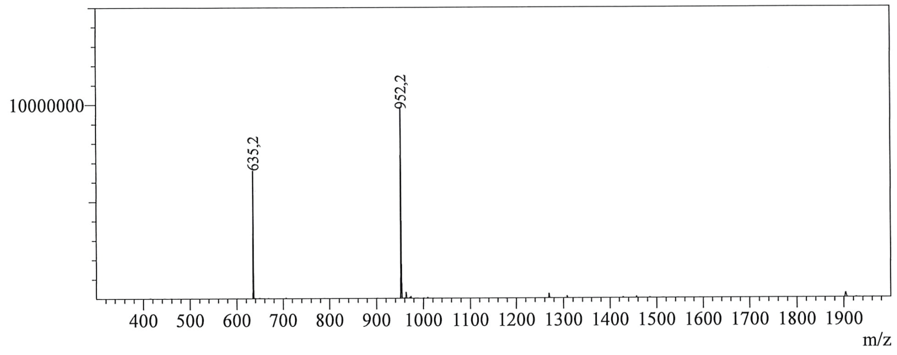


### Analog 6 - ESI-MS

Mass Spectrum
A69-14X1_28102019_652.lcd
Line#:1 R.Time:----(Scan#:----)
MassPeaks:1
Spectrum Mode:Averaged 0,107-0,207(33-63) BasePeak:888,1(7235119)
BG Mode:Averaged 0,493-0,700(149-211) Segment 1 - Event 1

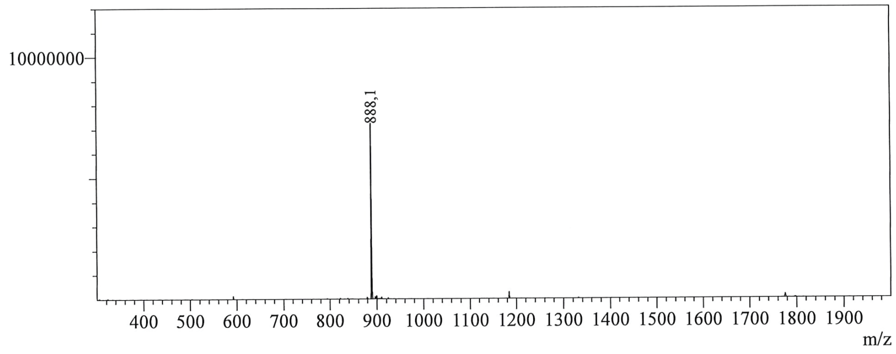
